# Mitochondrial Permeability Transition in Skeletal Muscle Phenocopies Muscle Alterations seen in Cancer Cachexia and other Wasting Conditions

**DOI:** 10.64898/2026.02.12.705530

**Authors:** Maya Semel, Cole Lukasiewicz, Sarah Skinner, Mark R. Viggars, Sung Gi Noh, Martin Picard, Abbey-Gale Mannings, Michael S. Cohen, Dennis W. Wolan, Terence E. Ryan, Russell T. Hepple

## Abstract

Skeletal muscle in wasting conditions often exhibits atrophy, mitochondrial respiratory dysfunction, and fragmentation of the acetylcholine receptor (AChR) cluster at the endplate. The accompanying alterations of mitochondrial morphology suggest mitochondria may be involved in muscle pathology in these conditions. To address this gap, we tested an established pathological mechanism in ischemia-reperfusion injury and neurodegeneration but poorly studied in skeletal muscle: mitochondrial permeability transition (mPT). We tested if mPT recapitulated phenotypes common in wasting conditions, whether tumor-conditioned media (TCM) could promote mPT, and compared differentially expressed genes (DEGs) induced by mPT with DEGs observed in a mouse model of pancreatic cancer cachexia. Inducing mPT in mouse skeletal muscle bundles progressively altered mitochondrial cristae morphology, culminating in breach of the outer mitochondrial membrane. Inducing mPT in mouse muscle fibers increased mROS and Caspase 3 activity and caused atrophy. Inducing mPT caused a complex I mitochondrial respiratory impairment, increased lysosome-mitochondrion co-localization, and fragmented the AChR cluster at the muscle endplate. The Ca^2+^ threshold for mPT, mitochondrial calcein colocalization and mitochondrial membrane potential were reduced by TCM in skeletal muscle or C2C12 myoblasts, respectively. Knockout of the mPT-regulating protein CypD attenuated the reduction in Ca^2+^ threshold for mPT by TCM. Inhibitors of mPT attenuated atrophy with TCM in C2C12 and human primary myotubes. Finally, there was overlap between the DEGs of mPT and diaphragm muscle in a mouse model of pancreatic cancer cachexia during the muscle wasting phase. We conclude that mPT should be explored as a therapeutic target in muscle wasting disorders.

**Graphic Abstract Legend:** Skeletal muscle wasting conditions are characterized by muscle fiber atrophy, mitochondrial respiratory dysfunction, mitochondrial depletion, and fragmentation of acetylcholine receptor (AChR) clusters at the neuromuscular junction. Here, we identify mitochondrial permeability transition (mPT) as a mechanism that recapitulates these pathological features. mPT was induced pharmacologically with Bz423 or Ferutinin, or promoted by tumor-derived factors present in tumor-conditioned media. Induction of mPT increased mitochondrial reactive oxygen species (mROS) production and Caspase-3 (Casp3) activation, resulting in muscle fiber atrophy and AChR fragmentation. Furthermore, mPT induction in C2C12 and human primary myotubes caused myotube atrophy, which was attenuated by pharmacological inhibition of mPT. mPT also caused a complex I-specific mitochondrial respiratory impairment and increased mitochondrial co-localization with lysosomes. Together, these findings identify mPT as a novel mechanism of pathological phenotypes associated with muscle wasting and support mPT as a promising therapeutic target.

**Key Points:**

- Mitochondrial permeability transition (mPT) induces marked alterations in mitochondrial morphology and muscle phenotypes that are common in wasting conditions
- mPT is promoted by tumor-derived factors in a manner that depends upon the mPT-regulating protein CypD
- mPT generates transcriptional alterations that overlap with cachectic muscle in a mouse model of pancreatic cancer, particularly during the period of muscle wasting
- Pharmacological targeting of mPT attenuates or prevents atrophy in C2C12 and human primary myotubes, respectively

## Introduction

Skeletal muscle in muscle wasting disorders frequently presents with a common set of phenotypes including muscle atrophy (Friedrich *et al*., 2015; Martin *et al*., 2023), mitochondrial respiratory dysfunction (Protti *et al*., 2007; Neyroud *et al*., 2019; Kunz *et al*., 2022), and dismantling of the acetylcholine receptor (AChR) cluster at the muscle endplate (Liu *et al*., 2016; Sartori *et al*., 2021). Furthermore, mitochondrial morphology is often found to be perturbed (Fontes-Oliveira *et al*., 2013; Owen *et al*., 2019; Zhang *et al*., 2023) and mitochondrial content reduced in these conditions (Brown *et al*., 2017; Neyroud *et al*., 2019). This has led to widespread conjecture that mitochondria are likely to be involved in driving muscle pathology (Gan *et al*., 2018b; De Mario *et al*., 2021), although the details remain unclear. It is notable that an event known as mitochondrial permeability transition (mPT) is well-known to mediate pathology in cardiac ischemia-reperfusion injury (Bertero *et al*., 2024), neurodegeneration (Godoy *et al*., 2021), and other contexts (Ma *et al*., 2021; Sautchuk *et al*., 2023), yet is scarcely studied in skeletal muscle. mPT involves the formation of a pore (the mPT pore or mPTP) across the inner mitochondrial membrane in response to elevated mitochondrial matrix Ca^2+^ (Bernardi *et al*., 2021). Prolonged mPT is associated with loss of proton motive force, inducing osmotic stress secondary to the H^+^ movement from the mitochondrial inter-membrane space into the matrix. This causes mitochondrial swelling, cristae displacement, and mitochondrial outer membrane rupture, releasing mitochondrion-localized factors into the cytoplasm to activate cell death, inflammatory and proteolytic pathways (Bernardi *et al*., 2015).

Therapeutic testing of targeting mPT to attenuate skeletal muscle pathology remains largely confined to muscular dystrophies (Millay *et al*., 2008; Reutenauer *et al*., 2008; Palma *et al*., 2009; Dubinin *et al*., 2020; Dubinin *et al*., 2021; Stocco *et al*., 2021), with a few studies in other conditions such as type II diabetes (Taddeo *et al*., 2014; Belosludtsev *et al*., 2021). Notwithstanding these studies, the role of mPT in driving skeletal muscle pathology remains obscure. To this end, we first established the impact of mPT on skeletal muscle mitochondrial morphology and its potential for promoting signals involved in activating muscle atrophy. We then assessed the impact of mPT on muscle fiber diameter (published previously in (Skinner *et al*., 2021)), mitochondrial respiratory function, mitochondrial co-localization with lysosomes (to gain insight to mitophagy), and integrity of the AChR cluster at the muscle endplate. To provide insight to the occurrence of mPT in muscle wasting conditions and its potential as a therapeutic target, we then determined whether tumor-derived factors associated with pancreatic cancer promote mPT in skeletal muscle, and whether this relied upon a key protein that regulates the Ca^2+^ threshold for inducing mPT, cyclophilin D (CypD). Finally, we tested for overlap in the transcriptional signature of mPT and that seen in a mouse model of pancreatic cancer cachexia, and the efficacy of pharmacological targeting of mPT to prevent atrophy induced by tumor-derived factors.

## Methods

### Ethical Approval

All mouse experiments were performed with approval of the respective animal care committees at the Children’s Hospital of Philadelphia and University of Florida (protocol number 202011171 and 202400000051). All mouse procedures were performed in compliance with *The Journal of Physiology’s* policies for ethical conduct of animal research.

### Animals

Experiments were performed using 3-6 month old male C57BL6J mice, and 4 month old male inducible muscle-specific CypD knockout (mkoPPIF) mice. Inducible mkoPPIF mice were generated by breeding PPIF^fl/fl^ mice (Jax stock no. 005737) with inducible muscle-specific Cre mice (Jax stock no. 031934). Cre was activated by *i.p.* Raloxifene injection (110 mg per kg given three times over the period of a week) and 4-6 weeks later muscles were used in incubation experiments to determine whether tumor-derived factors reduce the Ca^2+^ threshold for mPT. As evidence of the success of muscle-specific PPIF knockout in the mkoPPIF model, 4 weeks following raloxifene treatment there was negligible PPIF transcript and markedly reduced CypD protein in skeletal muscle of mkoPPIF mice, whereas liver levels were unaffected (Fig A1). Mice were housed in ventilated cages and provided food and water *ad libitum* and euthanized for terminal experiments by thoracotomy while under deep anesthesia with 2-4% isoflurane.

### Electron Microscopy of Mitochondrial Permeability Transition

To address the impact of mPT on skeletal muscle mitochondrial morphology, we induced mPT in C57BL6J mouse skeletal muscle mitochondria during a Ca^2+^ retention capacity (CRC) assay. Mice were deeply anesthetized using isoflurane (2-4%) before being euthanized by thoracotomy, and the gastrocnemius muscle was harvested, placed in ice-cold Buffer A (in mM: 2.77 CaK_2_EGTA, 7.23 K_2_EGTA, 6.56 MgCl_2_, 0.5 dithiothreitol (DTT), 50 K-MES, 20 imidazole, 20 taurine, 5.3 Na_2_ATP, 15 phosphocreatine, pH 7.3) and transferred to the stage of a light microscope for manual dissection into 3-4 mm length by 1 mm wide muscle bundles using sharp tweezers. Muscle bundles underwent a 30 min incubation in saponin (0.05 mg/ml)-supplemented Buffer A to permeabilize the muscle sarcolemmal membrane, followed by 3 washes in stabilizing Buffer B (in mM: 2.77 CaK_2_EGTA, 7.23 K_2_EGTA, 1.38 MgCl_2_, 3.0 K_2_HPO_4_, 0.5 dithiothreitol, 20 imidazole, 100 K-MES, 20 taurine, pH 7.3) and a phantomizing procedure to remove myosin as previously described (Picard *et al*., 2008). Bundles were kept in ice-cold CRC buffer (in mM: 250 sucrose, 10 MOPS, and 0.005 EGTA, 10 P_i_-Tris, pH 7.3) until use. Muscle bundles were placed in a cuvette of a fluorometer containing CRC buffer (37°C) with energizing substrates (5 mM glutamate, 2.5 mM malate) and Calcium Green 5N to permit monitoring of Ca^2+^ dynamics. As shown in Fig 1, muscle bundles were retrieved at 3 distinct phases of the Ca^2+^ uptake curve: (i) the rapid Ca^2+^ uptake phase (early); (ii) the point of reversal of the Ca^2+^ signal (middle); and (iii) the phase of mitochondrial Ca^2+^ release (end) and fixed for electron microscopy. Upon retrieval of the muscle bundle, it was placed in 2.5% (vol/vol) glutaraldehyde, 2% (vol/vol) paraformaldehyde in 0.1 M sodium cacodylate buffer (pH 7.4) at 4°C overnight. Bundles were washed, postfixed in 2.0% (wt/vol) osmium tetroxide at room temperature for 1 h, rinsed in distilled water and then stained with 2% (vol/vol) uranyl acetate. Bundles then underwent dehydration through a graded ethanol series and embedded in Embed-812 (Electron Microscopy Sciences). Thin sections were stained with uranyl acetate and lead citrate and imaged at 12,000x magnification on an electron microscope (JEOL 1010) using a Hamamatsu digital camera. A total of 18 bundles were used in these experiments. Analyses of mitochondrial morphology were made by a single observer blinded to the identity of the samples.

**Fig 1.**
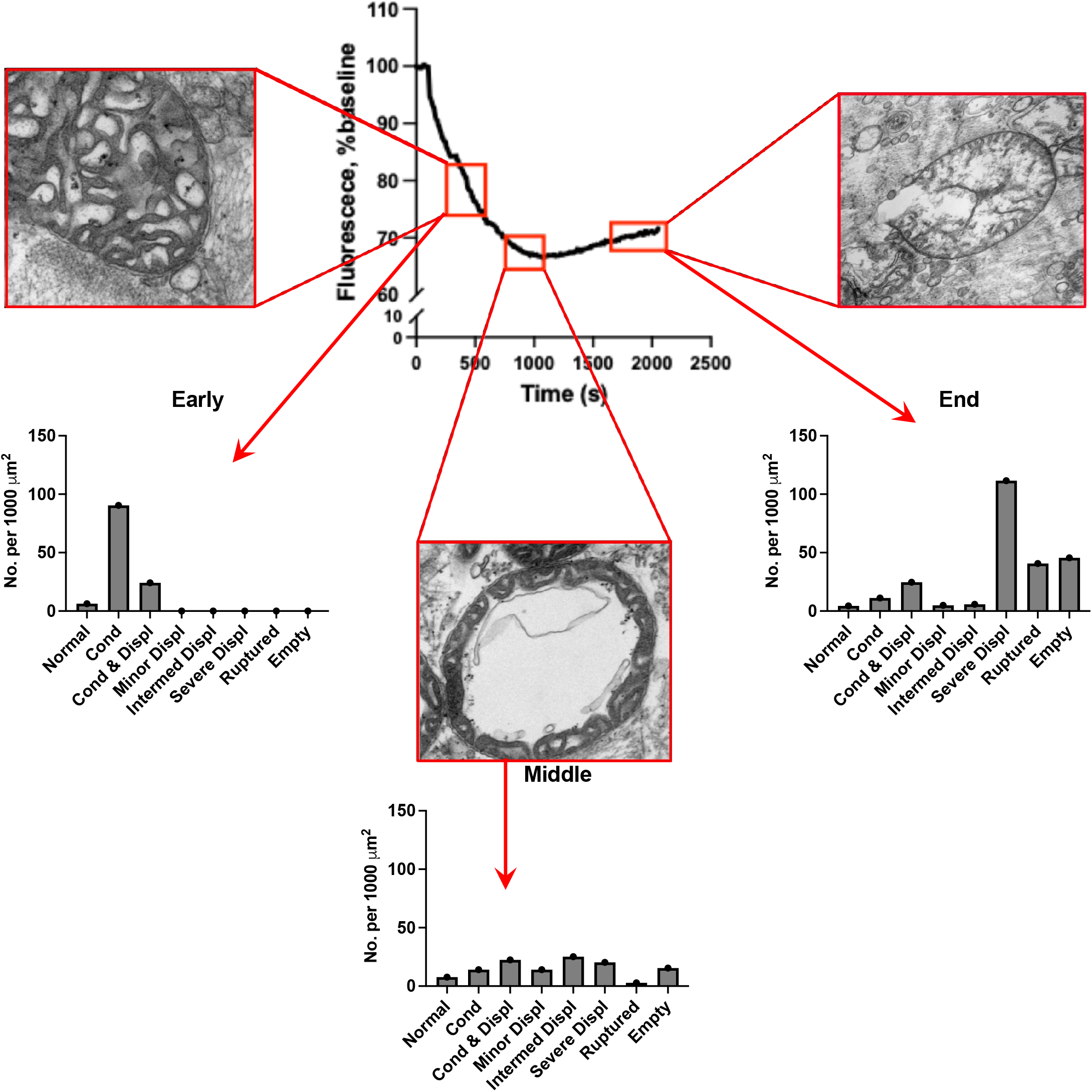
Morphological consequences of mPT for mitochondria. Skeletal muscle mitochondrial Ca^2+^ uptake that culminates in mPT causes progressive alterations in mitochondrial morphology, where the (i) initial phase is largely characterized by cristae condensation, with some cristae condensation accompanied by mild cristae displacement; (ii) the middle phase is associated with the appearance of mitochondria with varied levels of cristae displacement, some mitochondria with ruptured outer membranes, and others lacking discernible cristae; and (iii) the end phase characterized by large accumulation of mitochondria with severe cristae displacement, mitochondria with ruptured outer membranes and mitochondria lacking discernible cristae. Normal = normal morphology, Cond = condensed cristae, Cond & Displ = condensed cristae with cristae displacement, Minor Displ = minor cristae displacement, Intermed Displ = intermediate cristae displacement, Severe Displ = severe cristae displacement, Ruptured = ruptured outer mitochondrial membrane, and Empty = empty shell with no discernible cristae. Representative images at each phase show examples of mitochondria with cristae condensation (early), severe cristae displacement (middle), and rupture of the outer membrane (end). As this analysis was qualitative (no statistics), the values presented represent the mean across multiple microscope fields.

### Mitochondrial ROS, Caspase 3 FLICA, and Atrophy in Single Mouse Muscle Fibers

Data for these experiments has been published previously (Skinner *et al*., 2021) and is reproduced here to integrate with the bigger picture of how mPT affects skeletal muscle. As previously reported (Skinner *et al*., 2021), on the day of experiments C57BL6J mice were deeply anesthetized with isoflurane and the flexor digitorum brevis (FDB) muscles were removed and placed in room temperature physiological rodent saline. Mice were then euthanized by thoracotomy and removal of the heart. FDBs were incubated in a collagenase solution to isolate single fibers. Isolated single FDB fibers were then carefully transferred to 35 mm culture dishes (25-50 fibers per dish for 24 h experiments, and 50-100 fibers per dish for multi-day disuse experiments). For mitochondrial reactive oxygen species (mROS) experiments, single muscle fibers were incubated in media containing Vehicle, 300 nM Bz423 (to induce mPT), 300 nM Bz423 + 5 μM mitoTEMPO (to quench ROS), or 300 nM Bz423 + 1 μM TR002 (to inhibit mPT). For caspase 3 (Casp3) activity experiments, single muscle fibers were incubated in media containing Vehicle, 300 nM Bz423 (to induce mPT), 300 nM Bz423 + 20 μM Ac-ATS010-KE (to inhibit Casp3 activity), or 300 nM Bz423 + 1 μM TR002 (to inhibit mPT). For mROS experiments, muscle fibers were incubated for 30 min in MitoSOX (a live-cell permeable superoxide indicator). For Casp3 activity experiments, muscle fibers were incubated for 1 h with Casp3 FLICA reagent (a fluorescent probe targeted to activated Casp3). Muscle fibers were imaged at baseline and 24 h following treatments. Additional details of these approaches can be found in our previous publication (Skinner *et al*., 2021).

### Respiratory Function of Skeletal Muscle Mitochondria

To assess the impact of mPT on skeletal muscle mitochondrial respiration, we used high resolution respirometry to assess respiratory function in isolated mitochondria in the presence (50 μM Bz423, 25 and 50 μM ferutinin) or absence (vehicle, DMSO) of mPT induction. Mitochondrial isolation followed established protocols (Kim *et al*., 2023). Briefly, C57BL6J mice were anesthetized with isoflurane, and the gastrocnemius and quadriceps muscles were rapidly excised and placed in ice-cold dissection buffer (1x PBS, 1 mM EGTA). Visible blood, fat, and connective tissue were removed, followed by finely mincing the muscle. Minced muscle was transferred to a larger volume of dissection buffer supplemented with trypsin (0.0025%) and incubated on ice for 5 min to facilitate the release of intermyofibrillar mitochondria. Following incubation, the mixture was centrifuged at 200 x g for 5 min at 4°C, and the supernatant was aspirated and discarded.

The resulting pellet was resuspended in homogenization buffer (in mM, 50 MOPS, 100 KCl, 1 EGTA, 5 MgSO₄, 2 g/L bovine serum albumin; pH 7.1) and homogenized using a glass-Teflon homogenizer, then centrifuged at 800 x g for 10 min at 4°C to pellet nuclei and other non-mitochondrial components. The resulting supernatant was transferred to a new tube and centrifuged at 10,000 x g for 10 min at 4°C to pellet mitochondria. After aspiration of the supernatant, the pellet was gently washed in Resuspension buffer (in mM, 50 MOPS, 100 KCl, 1 EGTA, 5 MgSO₄; pH 7.1) to remove damaged mitochondria, then resuspended in resuspension buffer for respiration assays. Mitochondrial protein content was quantified using a BCA Assay kit (Pierce; A55865) to normalize assay values per µg of protein.

Mitochondrial respiratory function was assessed in isolated skeletal muscle mitochondria using an Oroboros O2k high-resolution respirometer (Oroboros Instruments, Innsbruck, Austria). A standardized volume of mitochondria was added to 2 mL of buffer D_r_ (in mM, 5 MgCl₂, 105 KCl, 10 KH₂PO₄, 5 MOPS, 1 EGTA, 2.5 g/L BSA; pH 7.2) supplemented with 5 mM creatine. Respiration assays were performed in the presence of vehicle (DMSO), Bz423 (50 µM), or ferutinin (25 and 50 μM) at 37°C under air-saturated oxygen conditions (∼200 µM O_2_). Following mitochondrial addition to the chamber containing Vehicle, Bz423 or ferutinin, oxygen consumption was recorded under basal conditions until a stable rate was achieved (∼3-5 min). Sequential substrate additions of glutamate (10 mM) and malate (2.5 mM), ADP (2 mM), and then succinate (10 mM) were used to assess state II, state III complex I-supported, and state III complex I+II-supported respiratory rates. Respiratory assays utilized 6 biological replicates (n=6 mice) for the Bz423 treatments and 4 biological replicates for the ferutinin treatments, with data from two chambers averaged per sample. In a subset of experiments (n=2 mice), we also added 10 μM Antimycin A (to inhibit Complex III), 10 mM ascorbate and 1 mM of the artificial electron donor *N,N,N’,N’-tetramethyl-p-phenylenediamine* (TMPD) following the Succinate step to test the capacity of Complex IV-driven respiration independent of complex I electron input to Complex III.

### Mitochondrial-lysosomal Colocalization in C2C12 Myotubes

C2C12 myoblasts were seeded in 8-well chambered slides (Ibidi) in DMEM with 10% FBS and 1% pen/strep for 48 h. Differentiation media (DM; DMEM, 2% horse serum, and 1% pen/strep) was added to the cells for four days and on day four MitoTracker Green (200 nM, Invitrogen) and LysoTracker Deep Red (500 nM, Invitrogen) dyes were added and incubated at 37°C for 15 min and 10 min, respectively. Baseline images were collected with a Leica SP8 Confocal Microscope (63x oil-immersion objective) and the following treatments were then added in duplicate: vehicle (0.2% DMSO), 1 μM FCCP, 5 μM Bz423, and 5 μM Bz423 + 10 μM TR002. Images were collected again 15 h post-treatment (based on pilot studies showing Bz423 had induced strong co-localization at this time point) and all images were analyzed for lysosome colocalization with mitochondria using the Huygens Essential software.

### AChR Cluster Integrity on Single Mouse Muscle Fibers

C57BL6J mice were deeply anesthetized using isoflurane and then underwent removal of the FDB muscle from both legs before being euthanized. The FDB muscles then were placed in a collagenase enzymatic solution to isolate single muscle fibers as we have done previously (Skinner *et al*., 2021). Muscle fibers were incubated in 2 mL of maintenance media containing AlexaFluor 488 conjugated α-bungarotoxin (1:1000) for labeling the AChRs. Approximately 8-10 fibers were transferred to a 35mm glass-bottom dish containing 2 mL of Dulbecco’s phosphate-buffered saline (DPBS) and DMSO, 300 nM Bz423 (to induce mPT), 300 nM Bz423 + 1 µM TR002 (to inhibit mPT), or 300 nM Bz423 + 20 µM Ac-ATS010-KE (to inhibit caspase 3). Muscle fibers were maintained under standard cell culture conditions (37°C, 5% CO_2_) for the duration of the experiments. Using a Leica SP8 confocal with 40x objective (total magnification 800x), 488 laser, and stage-top incubation system (Tokai Hit, Shizuoka, Japan), AChR clusters on each fiber were imaged using a small z-stack (<5 µm) at baseline and 12 h post-treatment. This time point was chosen as pilot experiments indicated that substantial changes occurred in the AChR cluster morphology by 12 h. Individual muscle fibers were tracked and matched for each timepoint to provide for repeat measurements of the same AChR clusters. All images were exported as maximal projections for analysis with NIH ImageJ. All AChR images were set to a threshold using the Huang method. Images used for AChR area were measured through automated selection of the α-bungarotoxin stained area. AChR fragmentation was also assessed using the ImageJ ‘Analyze Particles’ function to automatically identify the number of AChR clusters. The lower threshold limit was set to 0.5 µm^2^ as our group [32] and others [64] have done previously.

### Transcriptional Assessment of Bz423 induced mPT in Murine Myotubes

To explore the transcriptional alterations induced by mPT, C2C12 myoblasts (ATCC) at passage four were seeded in 6-well plates at 2.5×10^5^ cells per well and cultured for 2-days in growth media (10% FBS and 1% pen/strep). At ∼70% confluency, differentiation media (DM; 2% horse serum and 1% pen/strep) was added and refreshed every 48 h. Following 5-days in DM, myotubes were treated with either vehicle (0.1% DMSO) or 5 μM Bz423 for 12, 24, and 48 h in DM (n=6 technical replicates per condition/timepoint). At each timepoint, total RNA was isolated using TRIzol reagent (Invitrogen) followed by extraction with chloroform and precipitation with isopropanol. Following DNase treatment and removal using the DNA-free DNA removal kit (Invitrogen), high quality RNA samples were run on a high sensitivity RNA ScreenTape with Tapestation. Average RIN values were 9.23 ± 0.26. We used 1.25 µg of RNA as input for library preparation using Illumina mRNA-seq library Miniaturization with the Mosquito® robotic liquid handler by the University of Florida ICBR Gene Expression and Genotyping Core Facility, RRID:SCR_019145. Libraries were pooled with equal molarities and sequenced on a NovaSeq X by University of Florida ICBR NextGen DNA Sequencing Core Facility, RRID:SCR_019152. An average of 57 million 150bp paired-end reads were generated per sample. Adapter sequences were trimmed using fastp (v0.23.4). Paired end reads were aligned to mm10 using STAR (v2.7.11b) and gene expression quantification was performed using Salmon (v1.10.3) by University of Florida ICBR Bioinformatics Core Facility, RRID:SCR_019120. Quality control (QC) analysis was performed on raw count data after filtering genes with total counts ≥10 across all samples. Counts were normalized by library size (CPM) and log2(CPM+1) transformed for multivariate outlier detection. Principal component analysis (PCA) was applied, retaining the top components explaining >80% of variance. Robust Mahalanobis distances (MD²) were computed in PCA space using the Minimum Covariance Determinant estimator. MD² values were converted to p-values and adjusted for multiple testing using the Benjamini–Hochberg procedure; samples with q < 0.05 were flagged as potential outliers. Based on this criterion, one sample (Vehicle, 48 h, replicate 1) was removed prior to downstream analysis.

DESeq2’s negative binomial general linear model with dispersion estimation and Wald tests for contrasts was used (Love et al., 2014) to identify differentially expressed genes (DEGs) using a False Discovery Rate (FDR) of 0.05; the results of which are available within Supplementary Dataset 1. DAVID was used for pathway analysis using the background of all identified genes (total counts ≥10 across all samples) against KEGG, Reactome and GO Biological Processes databases. For pathway analysis presentation, pathway redundancy was reduced by filtering out pathways with more than 80% gene overlap within each list, keeping the pathway with the lowest FDR in each redundant group.

### Generation of Tumor Conditioned Media (TCM)

Murine pancreatic cancer (KPC 1245) cells, which are derived from the tumor of the KPC (*LSL-Kras*^G12D/+^; *LSL-Trp53*^R172H/+^; Pdx-1-Cre) mouse (Hingorani *et al*., 2005), were seeded and cultured overnight in DMEM with 1% pen/strep. Cells were then washed with 1X PBS, and serum-free media was added for 24 h. Media was collected and centrifuged at 2,500 rpm for 20 min, and the supernatant was filtered through a 0.22 μm filter and stored at −80°C for future use.

### Calcein-Cobalt Assay in C2C12 Myoblasts

To assess whether tumor-derived factors can induce mPT pore opening, murine C2C12 myoblasts were treated for 4 h with a vehicle (0.5% DMSO), KPC TCM (33%), or KPC TCM (33%) with the mPT inhibitor Alisporivir (5 µM, MCE). For the KPC TCM + Alisporivir group, cells were pre-treated with Alisporivir (5 µM) for 1 h prior to the addition of KPC TCM and remained in the presence of Alisporivir (5 µM) during the subsequent 4 h treatment. The cells were then assessed for quenching of mitochondrial calcein-AM fluorescence by cobalt in a calcein-cobalt mPT assay (Petronilli *et al*., 1999). Cells were coloaded with calcein-AM (1 µM, Invitrogen), MitoTracker Deep Red FM (150 nM, Invitrogen), and tetramethylrhodamine, methyl ester (TMRM; 50 nM, Invitrogen) at 37°C for 30 min, followed by CoCl_2_ (1 mM, Sigma) for another 30 min at 37°C, both in phenol red-free 1X HBSS (Gibco). Cells were then washed three times in 1X HBSS and live cell images (3 per well) were captured with a Leica SP8 Confocal Microscope using the 63x objective and the 488 nm (Calcein), 638 nm (MitoTracker), and 552 nm (TMRM) laser lines. Images were deconvolved and analyzed for colocalization of calcein with TMRM using Huygens Essential software. To analyze TMRM fluorescence intensity, a mask of the MitoTracker channel was generated and applied to the TMRM channel using ImageJ. The calcein-cobalt assay utilized two technical replicates per treatment within a plate for a single experiment and was repeated three times. To provide demonstration that ferutinin induces mPT (used in respirometry experiments, above), we performed a single experiment with C2C12 myoblasts with the calcein-cobalt assay as above, except that the incubation time upon addition of 12.5 μM ferutinin was only 30 min before loading the cells with calcein-AM.

### Identification of Common Gene Signatures Induced by In-vitro mPT and during the Time-course of Orthotopic KPC-induced Cachexia Development in the Murine Diaphragm

We extracted differentially expressed genes (DEGs) (FDR=0.05) from a publicly accessible RNA-sequencing dataset (PMCID: PMC11733308) from diaphragm muscles harvested from mice-bearing orthotopic murine pancreatic (KPC) tumors at time points reflective of pre-cachexia (D8 and D10), mild-moderate cachexia (D12 and D14) and advanced cachexia (endpoint). We compared the DEGs from this temporal transcriptomic dataset with the DEGs from 24 h post Bz423 treatment, which represents the largest magnitude of change in our *in-vitro* mPT model, stratified by directionality. We performed pathway analysis and pathway redundancy as described above, with data available in Supplementary Dataset 3.

### Ca^2+^ Retention Capacity Assay in Phantom Muscle Bundles

To assess the impact of tumor-conditioned media (TCM) on skeletal muscle sensitivity to mPT, skeletal muscle bundles were incubated in TCM derived from KPC pancreatic cancer cells. Male C57BL/6J, mkoPPIF, or PPIF^fl/fl^ mice were used to assess the impact of TCM on CRC with and without CypD inhibition (n=3 mice per genotype). The present protocol is adapted from previous work by our group (Gouspillou *et al*., 2014). The tibialis anterior (TA) was excised from anesthetized mice and immediately placed in a chilled petri dish containing Buffer A. The red region of the TA was mechanically dissected into 3-6 mg bundles, which were incubated on ice for 60 min in either 50% TCM (diluted in Buffer A) or vehicle control (50% DMEM diluted in Buffer A). Following incubation, bundles were permeabilized for 30 min on ice in Buffer A containing saponin (0.05 mg/mL) and subsequently washed for 5 min in Buffer C (in mM: 80 K-MES, 50 HEPES, 20 taurine, 0.5 DTT, 10 MgCl₂, 10 ATP; pH 7.3) to limit over-permeabilization. Bundles then underwent phantomization by incubation for 30 min in Buffer D (in mM: 800 KCl, 50 HEPES, 20 taurine, 0.5 DTT, 10 MgCl₂, 10 ATP; pH 7.3) to extract myosin, followed by a final 5 min wash in CRC Buffer (pH adjusted to 7.3 at 4°C to be used on ice only at this step).

For CRC measurements, 1 mL of CRC Buffer was supplemented with glutamate (5 mM), malate (5 mM), inorganic phosphate (1mM), oligomycin (0.5 nM), Ca²⁺ Green-5N (1 µM), EGTA (10 µM), and CaCl₂ (30 µM) in a glass cuvette and warmed to 37°C. A subset of experiments using phantomized muscle bundles from C57BL/6J mice utilized the CypD-dependent mPT inhibitor alisporivir (1 µM) in the final cuvette. Final bundle mass was recorded following washes, and the bundle was immediately added to the warmed and supplemented CRC buffer in a fluorometer (FluoroMax Plus-C; Horiba Scientific). Fluorescence was monitored with excitation and emission wavelengths set to 505 and 535 nm, respectively, to assess the dynamics of mitochondrial Ca²⁺ uptake and release. CRC was defined as the total Ca²⁺ sequestered by mitochondria before Ca²⁺ release, indicated by inversion of the fluorescence signal. CRC analyses were performed using custom-written code in Igor Pro software (Wavemetrics). All washing steps were conducted at 4°C, and CRC measurements were performed at 37°C. CRC values were expressed relative to post-wash bundle mass.

### Atrophy Assay in C2C12 and Human Primary Myotubes

C2C12 myoblasts (ATCC, US) were seeded at 0.5×10^5^ cells per well in a 24-well plate and cultured for 2 days in growth media (10% FBS and 1% pen/strep). At ∼70% confluency, differentiation media (DM; 2% horse serum and 1% pen/strep) was added and refreshed every 48 h. Following 5 days in DM, myotubes were treated with either vehicle (0.25% DMSO; n=4 wells), 12.5 µM ferutinin (n=5 wells), 33% KPC TCM (n=5 wells), 33% KPC TCM + 1 µM Alisporivir (Med Chem Express; n=5 wells), or 33% KPC TCM + 1 µM C105SR (Med Chem Express; n=5 wells) for 48 h in DM. All conditions were matched to the same final concentration of DMSO (0.25%). Brightfield images were collected on a Leica DMi8 microscope using a 10x objective and myotube width was measured at 5 locations along the length of each myotube in 30 myotubes per well using ImageJ.

Human primary muscle progenitor cells (MPCs; human myoblasts, P1) were plated in an uncoated flask in primary myoblast growth media (MPC GM; Ham’s F10, 20% FBS,1% pen/strep, and 5 ng/mL basic-FGF) and incubated at 37°C for 1 h and 15 min for fibroblast purification. The supernatant containing MPCs was transferred to a coated flask in MPC GM and allowed to proliferate to approximately 90% confluence. MPCs (P3) were seeded at a density of 1.2 x 10^5^ cells per well in a 48-well plate and differentiated into myotubes in MPC differentiation media (MPC DM; DMEM with 4.5 g/L glucose, 2% horse serum, 1% pen/strep, and 0.1% insulin/transferrin/selenium). After 6 days, cells were incubated with 33% KPC TCM, 33% serum free media (DMEM and 1% pen/strep; Control for serum depletion in KPC treatment), or 33% KPC TCM with one of: 0.1% DMSO (Vehicle for mPT inhibitors), 1μM C105SR, 1μM Alisporivir, or 1μM TR002 for 24 h. Subsequently, myotubes were washed twice with PBS and fixed/permeabilized in ice cold methanol/acetone (1:1) solution for 10 min at −20°C. Myotubes were incubated with a primary antibody against sarcomeric myosin heavy chain (MF 20, Developmental Studies Hybridoma Bank) at 1:25 in blocking solution (PBS, 5% normal goat serum, 1% BSA) for 1 h at 37°C. Myotubes were then incubated with a secondary antibody (AlexaFluor594, mouse IgG2b, ThermoFisher) at 1:250 for 1 h at 37° and imaged with a 10x objective on an EVOS FL Auto 2 imaging system. Four images per well in 6 replicates for each treatment were obtained. Myotube diameter was analyzed using the Myotube Analyzer (PMID: 35689270).

### Statistics

Data for the impact of mPT on mitochondrial morphology are presented in frequency histograms. Data for the impact of mPT on mitochondrial ROS, Caspase 3 activation, fiber size, and number of AChR cluster segments were analyzed using a nested One-way ANOVA with a Sidak post-hoc test. Data for the impact of mPT on mitochondrial respiration and co-localization of lysosomes with mitochondria were analyzed by One-way ANOVA with a Sidak post-hoc test, with the exception that state II respiration with Bz423 treatment was analyzed using a Wilcoxon matched-pairs signed rank test and state II respiration with 25 μM and 50 μM ferutinin treatment were analyzed by Student’s T-test. Data for the impact of KPC TCM on mitochondrial calcein co-localization, mitochondrial membrane potential (TMRM fluorescence), Ca^2+^ threshold for mPT, and myotube diameter were analyzed by One-way ANOVA with a Sidak post-hoc test. In all cases, the significance threshold was set as P<0.05. Data in figures show individual values with the bar height representing the mean and, where shown, the error bars indicate the standard deviation.

## Results

### mPT Causes Marked Alterations in Mitochondrial Morphology in Skeletal Muscle

We used phantom muscle bundles in a CRC assay and retrieved bundles at three distinct points in the Ca^2+^ uptake and release response: (i) the rapid Ca^2+^ uptake phase (early); (ii) the point of reversal of the Ca^2+^ signal (middle); and (iii) the phase of mitochondrial Ca^2+^ release (end). As summarized in Fig 1, mitochondria exhibited progressive changes in morphology, with the early phase of the Ca^2+^ uptake response largely characterized by condensation of the cristae, where some mitochondria with condensed cristae also exhibited minor displacement of the cristae (e.g., creating asymmetric gaps between cristae folds). In the middle phase mitochondria exhibited varying degrees of cristae displacement and occasionally exhibited a ruptured outer membrane or a complete lack of discernible cristae. In the end stage there was a large accumulation of mitochondria with severe cristae disruption, mitochondria with ruptured outer membranes, and mitochondria with no discernible cristae. Although not quantified in our analyses, we also observed mitochondria with outer membrane blebbing and accumulation of small vacuoles (some with probable cristae remnants) in the locations where mitochondria would normally be present (e.g., on either side of z-lines and subsarcolemmal region), that may suggest the mitochondria under the stress of high Ca^2+^ exposure were also undergoing fission and mitochondria-derived vesicle formation.

### mPT Increases Mitochondrial ROS Generation and Caspase 3 Activity and Reduces Diameter of Skeletal Muscle Fibers

We used a compound known as Bz423 that reportedly binds to the same proline residue on the oligomycin sensitivity conferring protein (OSCP) as CypD to promote mPT (Giorgio *et al*., 2013). Consistent with the known impact of Bz423 in promoting mPT, we found that 300 nM Bz423 reduced the Ca^2+^ threshold for mPT and this effect was ablated by a novel CypD-independent triazole mPT inhibitor called TR002 (1 μM; Fig 2A). As we have reported previously (Skinner *et al*., 2021) and shown here in Fig 2B and C, inducing mPT with 300 nM Bz423 increased both MitoSOX fluorescence (mitochondrial superoxide detector) and activated Caspase 3 fluorescence (Casp3 FLICA assay), and these effects could be prevented by either mPT inhibition with TR002 (1 μM), or by inhibiting mitochondrial ROS with mitoTEMPO (5 μM) or Casp3 with Ac-ATS1010-KE (20 μM), respectively. In addition, inducing mPT with Bz423 caused a significant reduction in muscle fiber diameter that was rescued by mPT inhibition (TR002), quenching mitochondrial ROS (mitoTEMPO) or inhibiting Casp3 activity (Ac-ATS0101-KE) (Fig 3).

**Fig 2.**
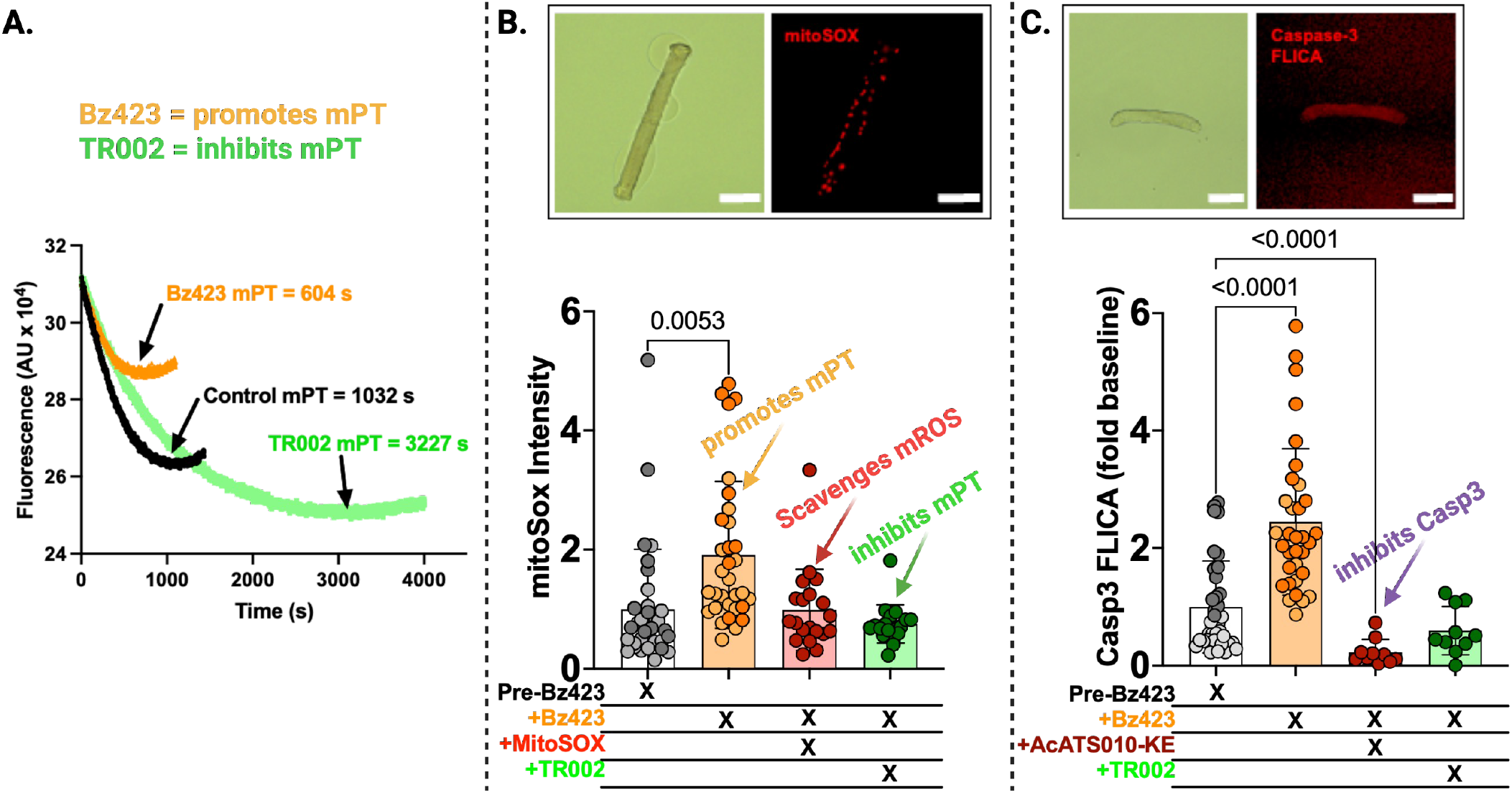
Impact of mitochondrial reactive oxygen species and caspase 3 activity in single mouse muscle fibers. Inducing mPT with 300 nM Bz423 in skeletal muscle fibers reduces the Ca^2+^ threshold for mPT, whereas treatment with the mPT inhibitor TR002 (1 ***μ***M) increases the Ca^2+^ threshold for mPT (A). Bz423-induced mPT promotes an increase in mitochondrial reactive oxygen species generation (B) and caspase 3 activity (C). One-way nested ANOVA with Sidak post-hoc test. Data in panels B&C were published previously in Skinner et al. Cells 2021.

**Fig 3.**
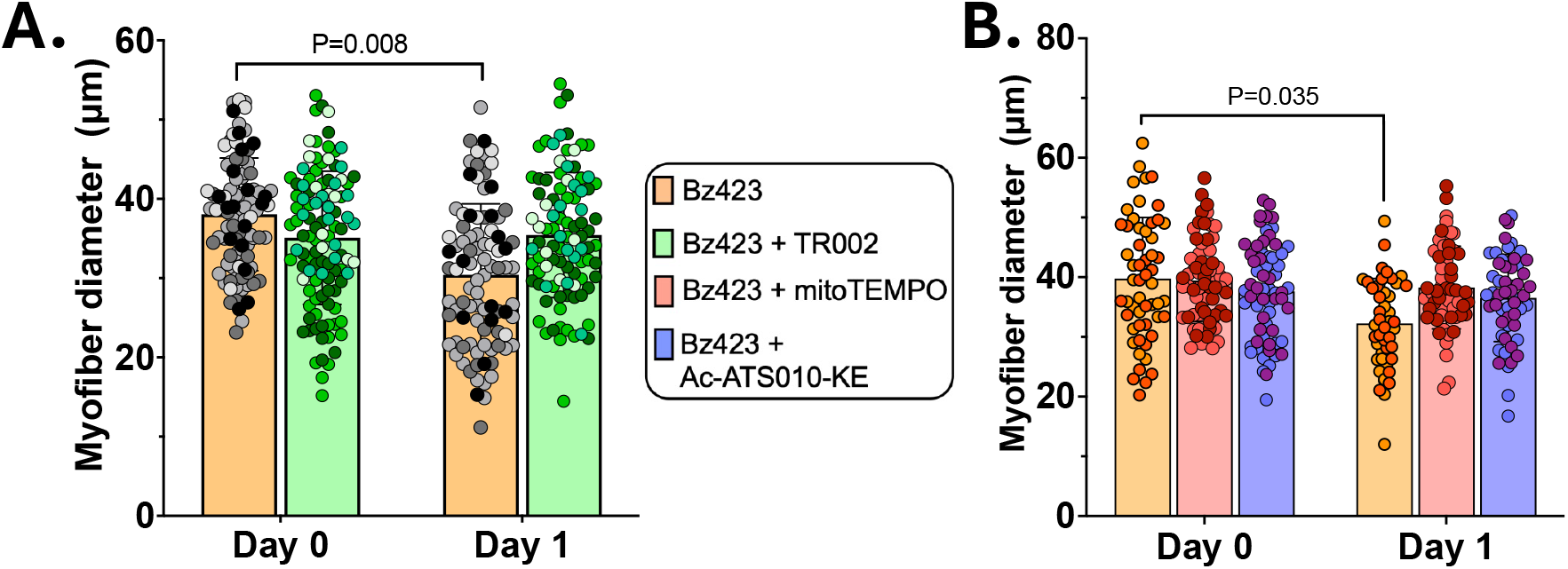
mPT induces muscle fiber atrophy. Inducing mPT (300 nM Bz423) in skeletal muscle causes atrophy (reduced fiber diameter) in single mouse flexor digitorum brevis muscle fibers, and this effect can be prevented by mPT inhibition (1 μM TR002) (panel A), quenching mitochondrial ROS (5 μM mitoTEMPO) or inhibiting Casp3 activity (20 μM Ac-ATS010-KE) (Panel B). One-way nested ANOVA with Sidak post-hoc test. Data were published previously in Skinner et al. Cells 2021.

### mPT Causes a Complex I-specific Impairment in Mitochondrial Respiratory Function and Increased Mitochondrial Co-localization with Lysosomes

Previously it was shown in isolated mitochondria from liver that inducing mPT by incremental additions of Ca^2+^ caused a complex I-specific deficit in mitochondrial respiratory capacity (Briston *et al*., 2017). To this end, we took mitochondria isolated from mouse hindlimb muscles and placed them in a polar electrode-based respirometer. We then assessed the changes in respiration in response to sequential addition of Complex I substrates (Glutamate, Malate), ADP, and a Complex II substrate (Succinate) in the presence of Bz423 (50 μM) or the Ca^2+^ ionophore ferutinin (25 μM or 50 μM) to induce mPT, or vehicle (DMSO). In a subset of experiments involving Bz423, we also added the artificial electron donor TMPD to provide electrons directly to complex IV to test if complex IV had a normal respiratory capacity with Bz423-induced mPT. Note that we provide in the appendix validation that ferutinin induces long-lasting mPT events based upon a reduction in mitochondrial calcein colocalization and reduction in mitochondrial membrane potential (TMRM) in a calcein-Cobalt assay (Fig A2).

Inducing mPT with Bz423 (Fig 4A & C) or ferutinin (25 μM in Fig 4B & D; 50 μM response shown in Fig A3) caused a dramatic reduction in the maximal rate of respiration with complex I substrates, whereas complex II-driven respiration was not different from vehicle. It was also noteworthy that state II respiration was elevated after inducing mPT with 50 μM Bz423 (P=0.03) and 25 μM (P=0.023) but not 50 μM ferutinin (Fig A4). Similarly, complex IV respiratory capacity, assessed after inhibition of Antimycin A and donating electrons directly to complex IV with TMPD, was unchanged following Bz423 treatment and yielded rates of respiration far in excess of the maximal rate of respiration seen with complex I and II substrates, substantiating the complex I-specific nature of this response. We also point out that this response is not compatible with an inhibition of ATP synthase by Bz423 because if that had occurred, we would have seen a reduced respiration with the TMPD condition.

**Fig 4.**
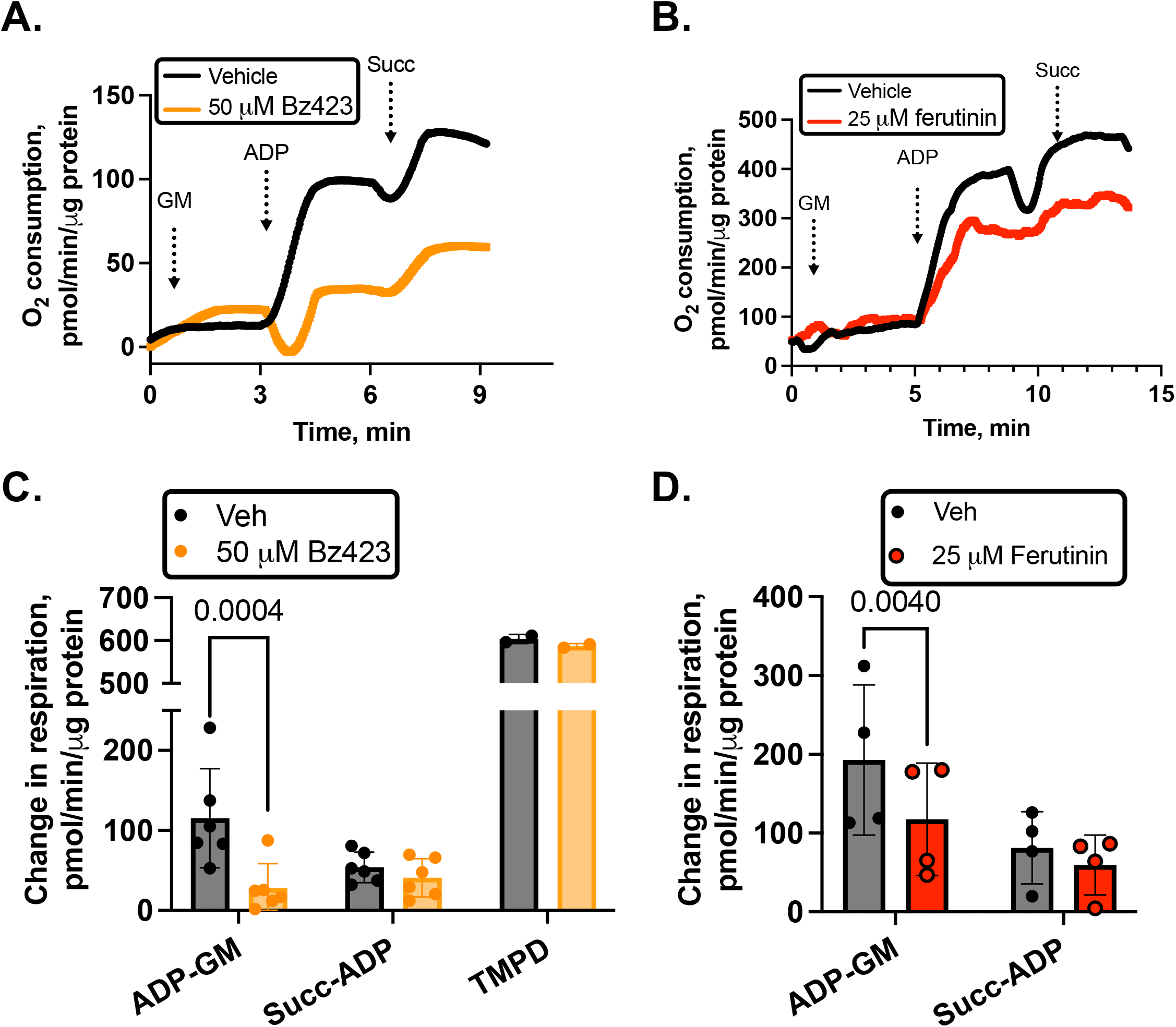
mPT causes a complex I-specific respiratory deficit in skeletal muscle mitochondria. A. Representative trace comparing the respiratory response to the complex I substrates glutamate-malate (GM), adenosine diphosphate (ADP), and the complex II substrate succinate (Succ) in Vehicle (black line) versus 50 μM Bz423 treated (orange) mitochondria. B. Representative trace for mPT induced using 25 μM the Ca^2+^ ionophore ferutinin (red) versus Vehicle (black) (same substrate conditions as in A). C. Summary data showing that whereas the increase in respiration upon addition of ADP with complex I substrates was markedly impaired in mitochondria treated with 50 μM Bz423 to induce mPT, the increase in respiration upon addition of the complex II substrate succinate was normal. Furthermore, in a subset of 2 mice we showed that the respiratory rate of complex IV following addition of TMPD was also unchanged by Bz423. D. Summary data for mPT induced by 25 μM ferutinin showing that similar to Bz423, mPT induced by ferutinin causes a complex I-specific respiratory deficit with a normal increase in respiration upon addition of the complex II substrate succinate. One-way ANOVA with Sidak post-hoc test.

To gain insight to the impact of mPT in stimulating mitophagy in skeletal muscle, we took C2C12 myotubes 5 days following transition to differentiation media and induced mPT in an immunofluorescent assay using MitoTracker Green and LysoTracker Deep Red. We induced mPT for 15h with 5 μM Bz423, and used the mitochondrial uncoupler, FCCP, as a positive control for the induction of mitophagy secondary to collapse of mitochondrial membrane potential. Representative confocal images are shown in Fig 5A. Consistent with the impact of mPT in promoting mitophagy in other cell types (Rodriguez-Enriquez *et al*., 2009; Song *et al*., 2015), inducing mPT with Bz423 in C2C12 myotubes increased the co-localization of mitochondria with lysosomes in a manner similar to the mitochondrial uncoupler FCCP. Furthermore, the impact of Bz423 was prevented by the mPT inhibitor TR002 (Fig 5B). We repeated this experiment on a separate day with freshly prepared reagents and obtained the same results (Fig A5).

**Fig 5.**
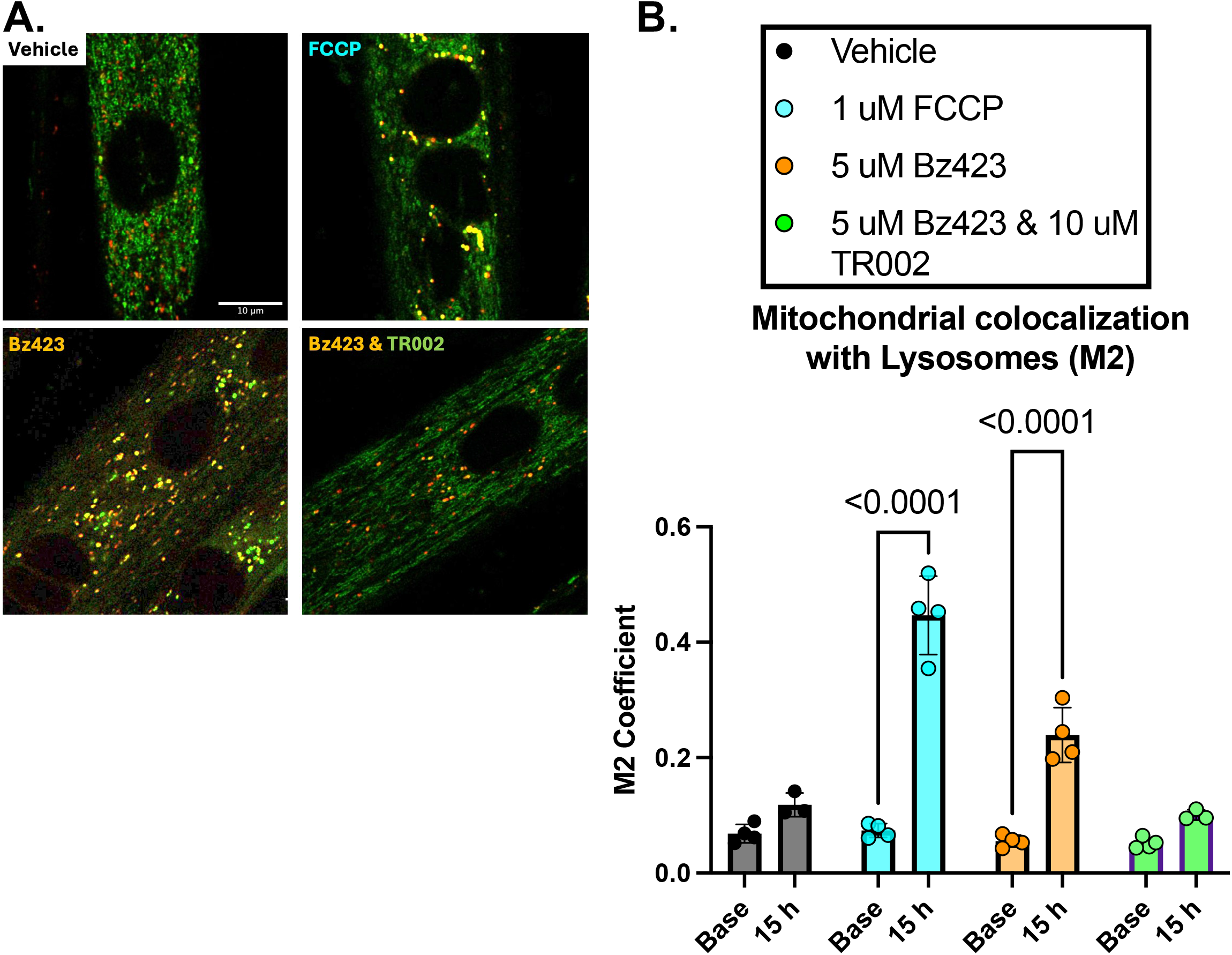
mPT increases lysosomal co-localization with mitochondria. A. Representative confocal images of myotubes where mitotracker are labeled in green and lysosomes are labeled in red. B. Summary results showing that inducing mPT with Bz423 increases the co-localization of lysosomes with mitochondria (M2 Mander’s Coefficient) and mPT inhibition with TR002 prevents this effect. Base = baseline measurements, 15 h = 15 hours post-treatment. Scale bar = 10 μm. One-way ANOVA with Sidak post-hoc test.

### mPT Causes AChR Cluster Fragmentation at the Muscle Endplate

A previous study identified that Casp3 dismantles the AChR cluster on C2C12 myotubes (Wang *et al*., 2014). Since our data show that mPT increases the activity of Casp3 in skeletal muscle (Fig 2B), we tested whether inducing mPT would cause an increase in the number of AChR cluster segments (a phenomenon known as AChR cluster fragmentation) on single mouse FDB muscle fibers. As shown in Fig 6, inducing mPT with Bz423 caused AChR cluster fragmentation, and this was prevented by either mPT inhibition (TR002) or Casp3 inhibition (Ac-ATS101-KE). A time-lapse video of this response is provided in an online Video (https://figshare.com/s/5c8cb696bfb639441ae8).

**Fig 6.**
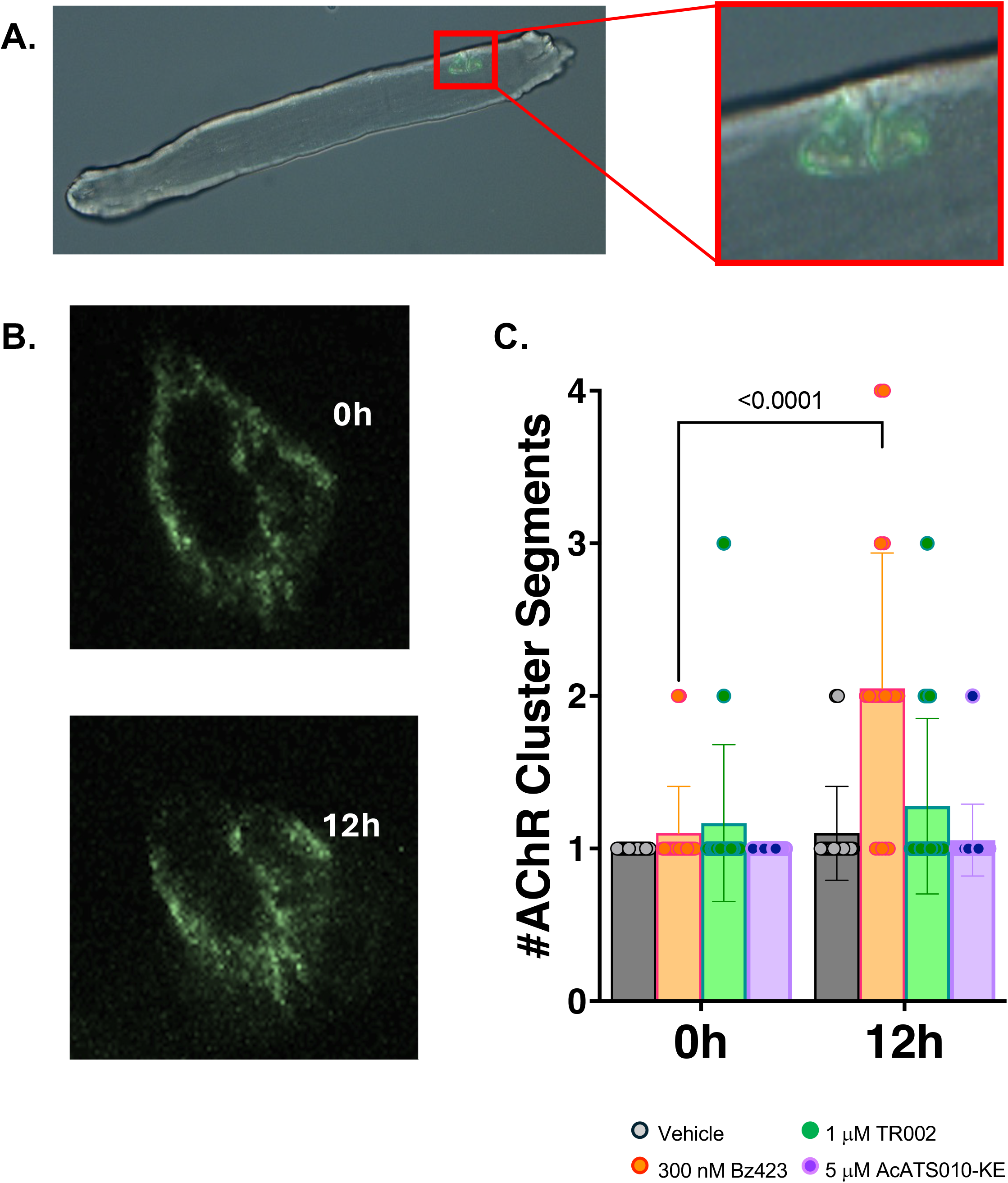
mPT causes dismantling of the acetylcholine receptor cluster at the muscle endplate. Panel A shows a single FDB muscle fiber labeled with AF488-fluorescently conjugated *α*-bungarotoxin (green). Panel B shows representative images of a single AChR cluster on a muscle fiber at baseline (0 h) and 12 hours (12 h) after treatment with 300 nM Bz423 to induce mPT. Bz423-induced AChR cluster fragmentation (indicated by an increased number of AChR segments) was prevented by mPT inhibition (1 μM TR002) or Casp3 inhibition (20 μM AcATS010-KE) (C). Nested One-way ANOVA with Sidak post-hoc test.

### Inducing mPT Alters the Muscle Transcriptome

To provide insights to the pathways downstream of mPT we used Bz423 to induce mPT in C2C12 myotubes and isolated RNA at 12, 24 and 48 h post-treatment for bulk RNA seq analysis. As shown in the principal component analysis in Fig 7A, Bz423-induced mPT generated widespread alterations in the transcriptome and this evolved considerably over the three time points including progressive upregulation of *Ucp2* and *Ucp3*, indicative of increased mitochondrial proton leak and oxidative stress concomitant with progressive downregulation of the adult AChR subunit (*Chrne*) that is lost with denervation (Fig 6B). A summary of the total number of differential expressed genes is presented in the Upset Plot, Fig 7C.

**Fig 7.**
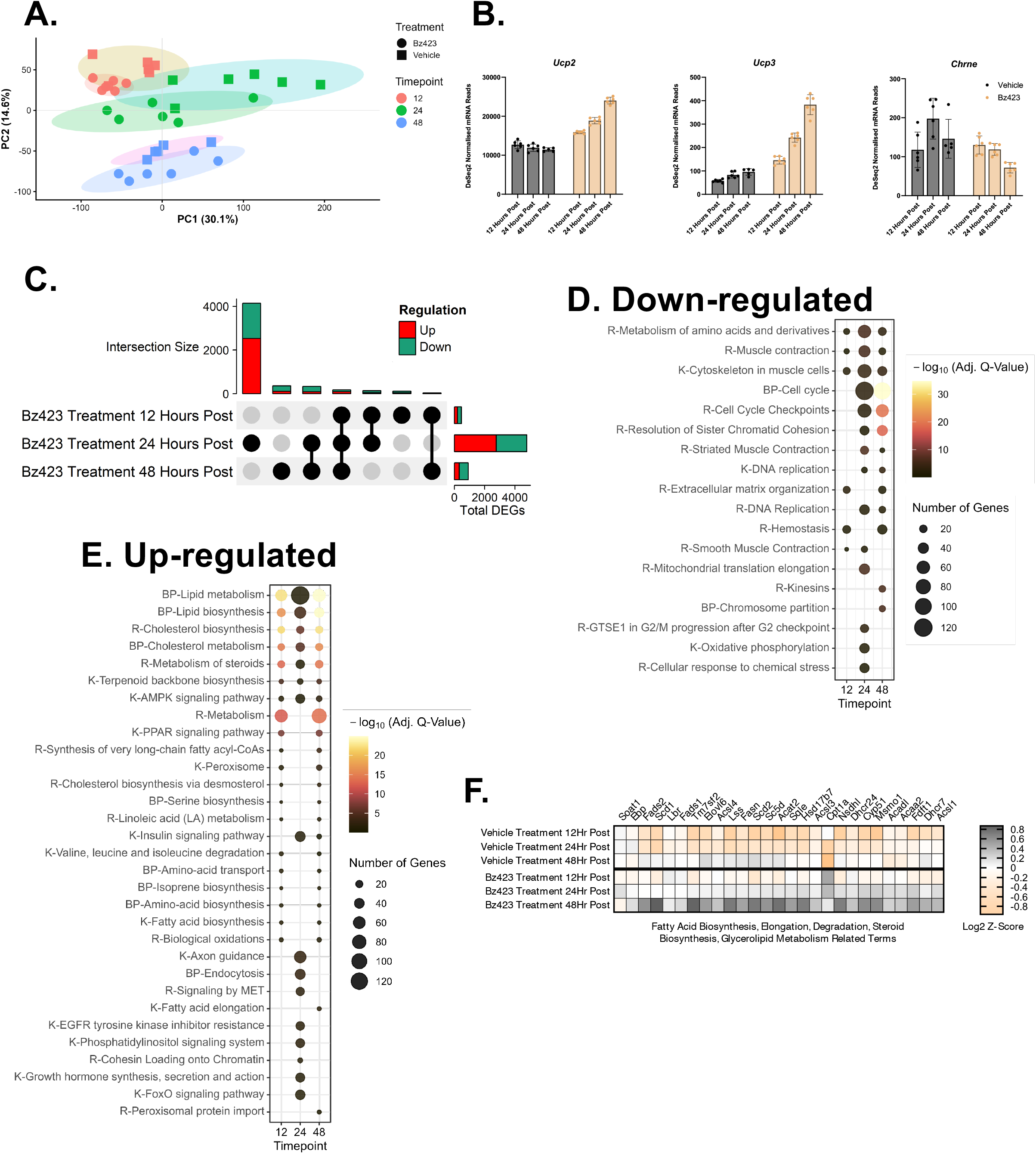
Transcriptional Signature of mPT. C2C12 myotubes (5 d post-differentiation) were incubated in 5 μM Bz423 to induce mPT. RNA was isolated at 12 h, 24 h and 48 h following treatment for RNAseq transcriptional profiling. A: Principal component analysis plot; B: Examples of genes increasing (Ucp2, Ucp3) or decreasing (Chrne) with mPT, C: Numbers of transcripts up- versus down-regulated over time following inducing mPT. D: Functional classifications of genes down-regulated by mPT; E: Functional classifications of genes up-regulated by mPT. F: Heat map showing alterations in fatty acid genes in response to mPT. K = Kegg, BP = Biological Processes, R = Reactome.

A total of 465 genes were differentially expressed (200- Up/265 Down) at the early timepoint (12h post-treatment). The early upregulated genes were dominated by pathways involved in fatty acid/lipid metabolism, perhaps in response to the mitochondrial membrane damage induced by mPT (Figure 7D). This included upregulation of the rate limiting enzyme of cholesterol biosynthesis (*Hmgcr*) and master regulator of *de novo* lipogenesis (*Fasn*). Additionally, several mitochondrial lipid-metabolism genes were upregulated, including *Cpt1a*, which mediates long-chain fatty-acid entry into mitochondria, and *Acsl1*, which activates fatty acids and directs them toward oxidative pathways. Upregulation of *Acadl*, the dehydrogenase initiating long-chain β-oxidation, and *Mlycd*, which reduces malonyl-CoA to relieve inhibition of *Cpt1a*, suggest early adjustments in mitochondrial fatty-acid oxidation capacity following mPT. Upregulation of *Elovl6*, which remodels C16–C18 fatty-acid chain length and influences mitochondrial membrane composition, further indicates remodeling of organelle lipid architecture in response to mPT. At the 12h post-treatment time point we also saw down- regulation of pathways in muscle cytoskeleton assembly and amino acid metabolism related genes including *Aldh4a1*, *Aldh7a1*, *Bcat2* and *Mccc2.* This suggests a mechanism where mitochondrial amino-acid catabolism and anaplerosis is downregulated to reduce carbon influx into the TCA cycle while oxidative phosphorylation is compromised by mPT (Fig 7E).

The most expansive changes in transcriptome signature were seen 24h post-treatment (4785 DEGs, 2763 Up, 2022 Down), where there were further increases in representation of fatty acid/lipid metabolism pathways (Fig 7F), as well as upregulation of 38 FoxO signaling-related genes, consistent with the atrophy that we observe with mPT (Skinner *et al*., 2021). We also observed at the 24h time point a further reduction in amino acid metabolic pathways, and reductions in pathways such as cell cycle, mitochondrial translation elongation, and oxidative phosphorylation genes. Changes 48h post-treatment were more modest and although some pathways exhibited greater log-fold changes, the number of genes within a given pathway was often reduced relative to 24h, (910 DEGs, 312 Up, 598 Down). Fig A5 shows targeted pathway heatmaps to highlight pathways related to the pathological phenotypes observed in muscle with inducing mPT. These changes were often small in magnitude (log2-fold changes ≤±0.2 for atrophy pathways in Fig A6A, mitochondrial oxidative phosphorylation genes in Fig A6C and autophagy/mitophagy genes in Fig A6D; log2-fold changes ≤±0.6 for neuromuscular junction genes in Fig A6B) but were consistent with the phenotypes observed.

### Tumor-Derived Factors Promote mPT in Skeletal Muscle

To address the potential for mPT in skeletal muscle to be promoted in cancer, we incubated C2C12 myoblasts in KPC TCM for 4 h and performed a calcein-cobalt assay. As shown in Figure 8A and B, KPC TCM reduced colocalization of calcein within mitochondria and reduced mitochondrial membrane potential, indicating that tumor-derived factors in KPC TCM directly induce long-lasting mPTP opening. Furthermore, co-treatment with the CypD-dependent mPT inhibitor Alisporivir attenuated these changes. We then incubated muscle fiber bundles in 50% KPC TCM for 60 min prior to saponin permeabilization and myosin depletion and then performed a CRC assay. As shown in Fig 8C, KPC TCM dramatically reduced the Ca^2+^ threshold for mPT and this was prevented by the CypD-dependent mPT inhibitor Alisporivir. Similarly, muscle-specific knockout of CypD attenuated the reduction in Ca^2+^ threshold induced by tumor-conditioned media by 75% if considered as a percentage of that seen in wildtype and by 50% if considered in absolute terms (8D). This shows that tumor-derived factors can promote mPT, in part by reducing the Ca^2+^ threshold for mPT in a manner that to a significant extent depends upon CypD.

**Fig 8.**
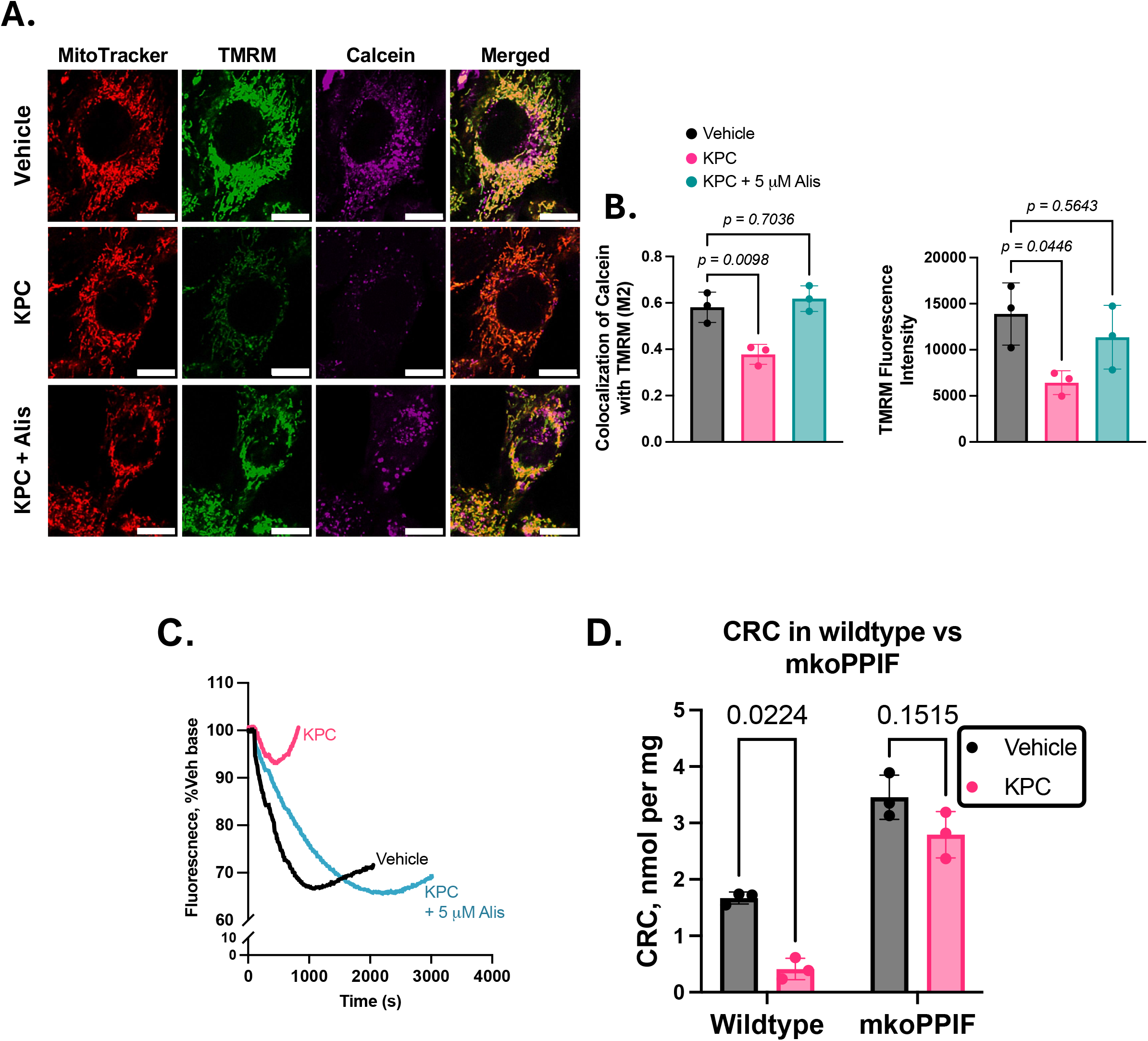
Tumor-conditioned media promotes mPT by reducing the Ca^2+^ threshold for mPT in a manner that depends upon CypD. We used a calcein-cobalt assay in C2C12 myoblasts where myoblasts were incubated in 33% KPC tumor-conditioned media for 4 h to test for impact on mPT. Representative images of the calcein, mitotracker (to label mitochondria), TMRM (mitochondrial membrane potential) and merged images of this assay are shown in panel A (scale bars = 10 µm). Tumor-conditioned media reduced mitochondrial Calcein and mitochondrial membrane potential, consistent with long-lasting mPT events, and this was rescued by mPT inhibition with Alisoporivir (Alis) (Panel B; n=3 independent experimental replicates). Panel C shows representative traces of the Ca^2+^ uptake and release response of muscle bundles incubated for 60 min in 50% KPC tumor-conditioned media. KPC tumor-conditioned media reduced the Ca^2+^ threshold for mPT in a manner that was largely dependent upon the mPT-regulating protein CypD (Panel D; n=3 mice per genotype with 2 bundles averaged per mouse in each treatment). In all cases the Vehicle condition contained an equal concentration of DMSO as the Alisporivir treatment. One way ANOVA with Sidak post-hoc test.

### Significant Overlap in Differentially Expressed Genes between Bz423-induced mPT and Skeletal Muscle from a Mouse Model of Pancreatic Cancer Cachexia

To gain further insight into the potential occurrence of mPT in cancer cachexia, we determined the extent of overlap between DEGs at the 24h timepoint of Bz423-induced mPT in C2C12 myotubes and a publicly accessible time-course RNA sequencing generated from diaphragm muscle of mice bearing orthotopic pancreatic KPC tumors (Neyroud *et al*., 2025). We saw significant overlap in DEGs in both directions that increased in magnitude over time following inoculation with pancreatic tumor cells when using a false discovery rate of 5% (Fig 9A). Notably, the overlap became particularly striking from day 12 to endpoint, which is the period in this model when diaphragm muscle wasting occurs. Further to this point, a plot of the diaphragm diameter (estimated from Fig 2F in (Neyroud *et al*., 2025)) versus the number of mPT-induced DEGs represented in the diaphragm transcriptome of mice with pancreatic cancer revealed a strong inverse relationship (Fig 9B). Summaries of the pathway analysis for up-regulated and down-regulated DEGs are shown in Fig 9C and 9D, respectively. Amongst the shared down-regulated DEGs between mPT and muscle in mice with pancreatic cancer were metabolic pathways (mitochondrial oxidative phosphorylation genes, TCA cycle), and protein synthesis pathways (ribosomes, protein translation and biosynthesis). Amongst the shared up-regulated DEGs were genes involved in lipid metabolism (cholesterol biosynthesis, sphingolipid signaling, and steroid biosynthesis pathways), neuromuscular junction remodeling (axon guidance and neurogenesis pathways), proteostasis related terms (protein digestion and absorption, ubiquitin conjugation pathways), and cell death (p53 signaling pathway). This analysis is consistent with mPT occurring in cancer cachexia during the period when muscle pathology is progressing and being an important contributor to pathways that are implicated in cancer cachexia muscle pathology.

**Fig 9.**
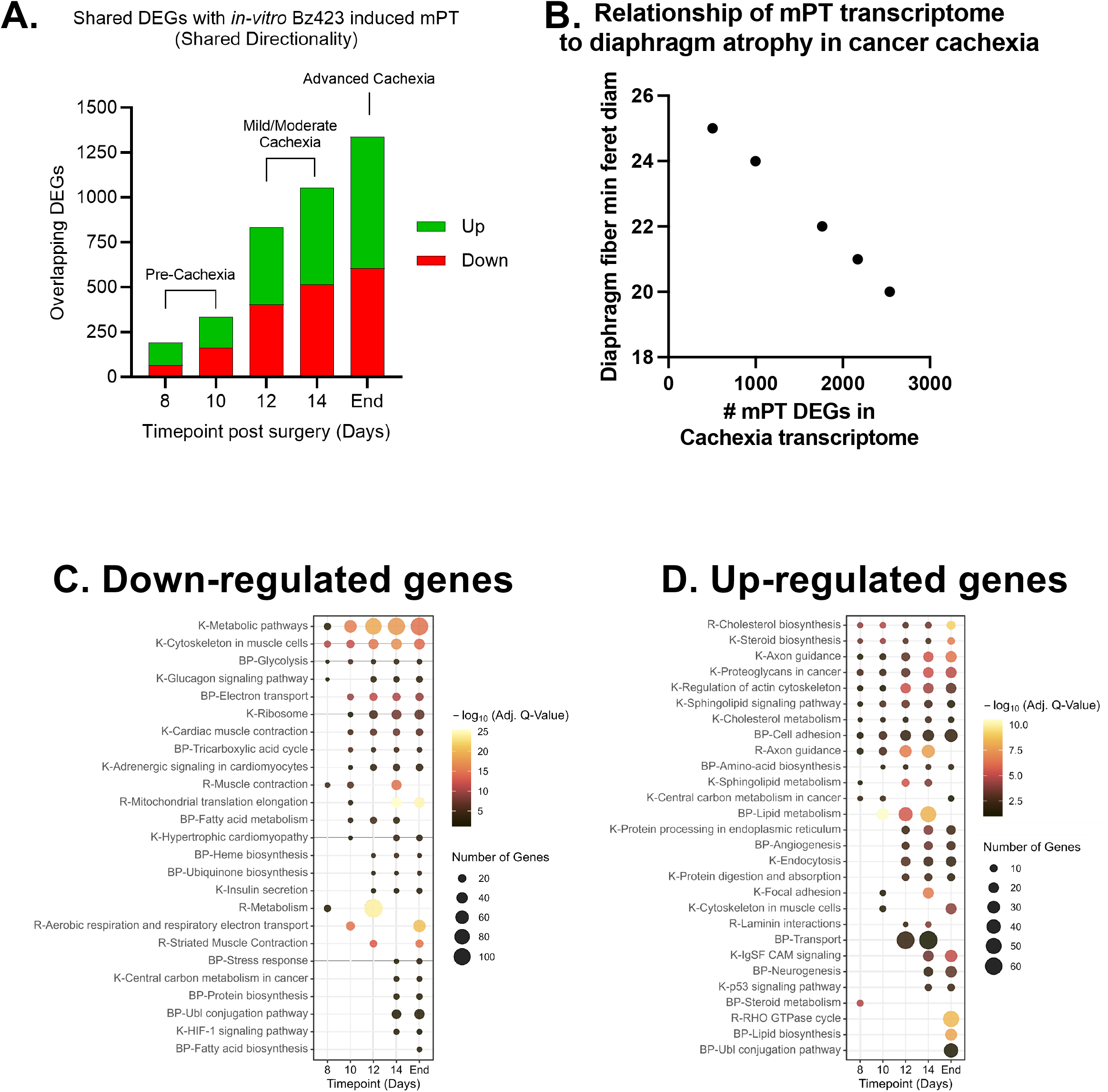
Overlap of differentially expressed genes (DEGs) between mPT *in vitro* and a mouse model of cancer cachexia. There was a progressive increase in the number of common DEGs between mPT (using the 24 h time-point) and in diaphragm muscle of mice with pancreatic cancer published previously (Neyroud et al. J Cachex Sarcopenia Muscle 16[1]: e13668, 2025), and this was particularly evident during the period of muscle wasting (A). The decline of diaphragm fiber size tracked an increase in mPT DEGs in the diaphragm transcriptome (B). Analysis of common DEGs with mPT and pancreatic cancer cachexia that are down-regulated (D) or up-regulated (E), with pathway analysis done by KEGG (K), Reactome (R), or (BP) (letters before pathway denote source).

**Fig 10.**
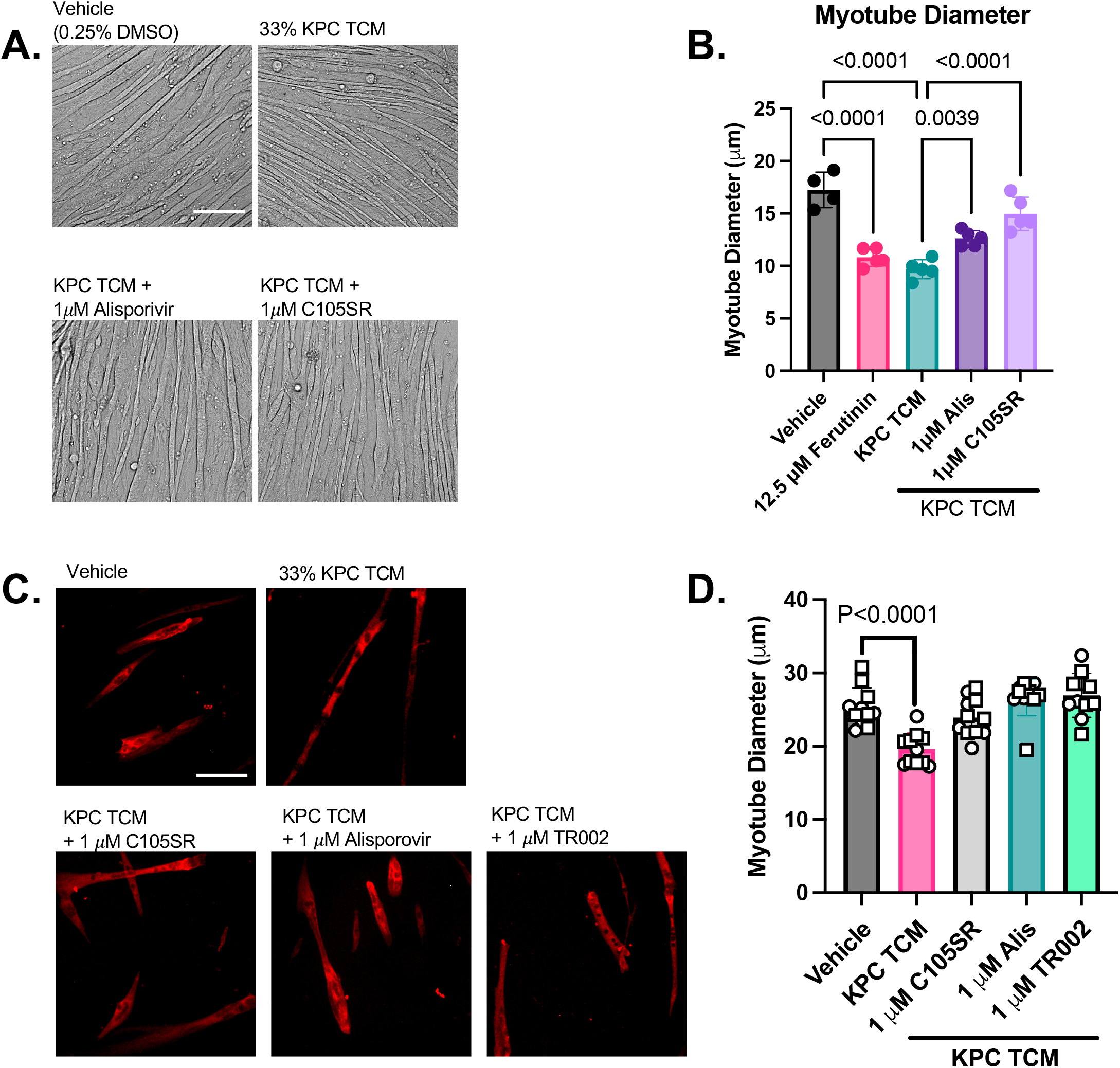
Inhibiting mPT attenuates atrophy induced by pancreatic tumor conditioned media in C2C12 and human primary myotubes. Representative images of C2C12 myotubes in each condition are shown in Panel A. Whereas 48 h incubation in KPC tumor conditioned media (TCM) induced marked myotube atrophy, two CypD-dependent mPT inhibitors, Alisporovir and C105SR, significantly reduced atrophy induced by KPC TCM (one independent experiment with n=4-5 wells per treatment; One way ANOVA with Sidak post-hoc test). The Ca^2+^ ionophore ferutinin was used a positive control for the impact of mPT on myotube diameter (Panel B). Representative images of human primary myotubes in each condition are shown in Panel C. Whereas KPC TCM induced significant myotube atrophy, both CypD-dependent (Alisporovir, C105SR) and independent (TR002) mPT inhibitors prevented myotube atrophy induced by KPC TCM (Panel D; two independent experiments differentiated by symbol shape, with n=5-6 wells per treatment in each experiment. One way ANOVA with Sidak post-hoc test). Scale bars = 100 µm.

### KPC Tumor Conditioned Media Causes Myotube Atrophy that is Attenuated by mPT Inhibition

To test the dependence of atrophy induced by tumor-derived factors on mPT we incubated C2C12 and human primary myotubes in KPC TCM for 48 h, with and without CypD-dependent (Alisporivir, C105SR) and independent (TR002; human primary myotubes only) mPT inhibitors. KPC TCM reduced myotube diameter relative to vehicle by 37% and 24% in C2C12 and human primary myotubes, respectively. In C2C12 myotubes although both mPT treated groups still exhibited atrophy relative to vehicle (Alisporivir reduced the atrophy to 23% and C105SR reduced the atrophy to 13%), myotubes in both mPT treated groups had significantly greater diameter than the KPC condition. The mPT inhibition was even more successful in human primary myotubes where none of the mPT treated groups exhibited significant atrophy relative to the vehicle control. This provides strong support for the idea that mPT is an important driver of atrophy in response to tumor-derived factors.

## Discussion

mPT is a well-established mechanism of tissue pathology in ischemia-reperfusion injury (Duchen *et al*., 1993; Baines *et al*., 2005; Theruvath *et al*., 2008), neurodegeneration (Savino *et al*., 2013), and a variety of other contexts such as inflammatory osteogenic dysfunction (Gan *et al*., 2018a) and inflammatory chondrocyte damage (Ma *et al*., 2021). Unfortunately, studies of mPT have yet to be performed in the majority conditions where alterations in mitochondrial biology and function may play a role in muscle pathology, such as cancer cachexia and other muscle wasting conditions. Furthermore, there is poor understanding of the consequences of mPT for skeletal muscle, which limits our ability to consider the full scope of its potential as a therapeutic target. To address these gaps, we performed a broad assessment of the consequences of mPT in skeletal muscle, focusing on altered muscle phenotypes that are common in wasting conditions. To address the potential relevance of mPT in cancer cachexia, a wasting condition where mitochondrial impact is well-established, we tested whether conditioned media from pancreatic tumor cells could reduce the Ca^2+^ threshold for mPT and tested its dependence on CypD, and tested whether atrophy induced by TCM could be attenuated by mPT inhibition in myotube cultures. Our findings reveal that mPT generates numerous phenotypes that are commonly seen in wasting conditions, including altered mitochondrial morphology that particularly impacts the cristae, increased ROS and Casp3 activity, muscle fiber atrophy, a complex I-specific respiratory impairment, increased mitochondrial co-localization with lysosomes, and fragmentation of the AChR cluster at the muscle endplate. Furthermore, we found that KPC TCM was sufficient to induce long-lasting mPT events and reduced the Ca^2+^ threshold for mPT in both C2C12 myoblasts and skeletal muscle fibers, respectively, in a manner that depended in part upon CypD. We observed marked overlap between the transcriptional alterations induced by mPT and those seen in diaphragm skeletal muscle of a mouse model of pancreatic cancer, particularly during the period when muscle wasting was occurring. Not only did mPT recapitulate many DEGs that are implicated in driving muscle pathology with cachexia, but there was also a striking proportionality between the number of mPT DEGs found in the diaphragm muscle transcriptome and the progression of diaphragm muscle atrophy in a mouse model of cancer cachexia. Collectively, our results suggest mPT should be explored for its role in driving muscle pathology in wasting conditions and evaluated as a therapeutic target to protect skeletal muscle in these states.

### History of mPT in Skeletal Muscle Research

Compared to the well-studied role of mPT in ischemia-reperfusion injury of the heart (Halestrap *et al*., 1998; Baines *et al*., 2005; Robichaux *et al*., 2023), mPT is not well studied in skeletal muscle pathology. Notwithstanding, Wrogeman and Pena proposed in 1976 that necrosis in muscular dystrophies was likely driven by Ca^2+^ overload of the mitochondria and that targeting mPT may offer a therapeutic approach that would be widely applicable to muscle conditions associated with Ca^2+^ dyshomeostasis (Wrogemann & Pena, 1976). The first direct evidence in support of this view came from studies published in 2008 and 2009 showing that genetic knockout of the mPT promoting protein CypD protected against muscle pathology in mouse models of muscular dystrophies that included delta sarcoglycan deficiency and Lama2 deficiency (Millay *et al*., 2008) and collagen VI deficiency (Palma *et al*., 2009). Since then, additional studies have shown that genetic approaches (Millay *et al*., 2008; Bround *et al*., 2023) or pharmacological approaches to reduce mPT (Reutenauer *et al*., 2008; Dubinin *et al*., 2021; Stocco *et al*., 2021) provide protection to skeletal muscle in animal models of different muscular dystrophies. Emerging areas where mPT is being considered in skeletal muscle pathology include diabetes (Belosludtsev *et al*., 2021) and nemaline myopathy (Gartz *et al*., 2023). However, in any of these scenarios the specific consequences of mPT are largely unknown, and the role of mPT in muscle wasting conditions has not yet been addressed.

### Mitochondrial Morphology in Wasting Disorders

Alterations in mitochondrial morphology are commonly reported in a wide range of wasting conditions, including cancer cachexia (Fontes-Oliveira *et al*., 2013; Zhang *et al*., 2023), sepsis (Owen *et al*., 2019; Pierre *et al*., 2025), and chronic kidney disease (Kouidi *et al*., 1998). Visually, the mitochondrial alterations bear resemblance to features seen in cells exposed to chemicals that trigger mitochondrial-mediated apoptosis (Sesso *et al*., 2004), suggesting that mPT may be involved. To directly address how mPT affects mitochondrial morphology in skeletal muscle, we retrieved muscle bundles in a CRC assay at three phases of the Ca^2+^ uptake and release response (early, middle, end). These experiments revealed progressive changes in mitochondrial morphology, first characterized by cristae condensation during the Ca^2+^ uptake phase (early), and progressing to varying degrees of cristae displacement in the middle and end phase, with a marked accumulation of mitochondria with severe cristae displacement and mitochondria with a ruptured outer membrane during the Ca^2+^ release phase (end). One example from the muscle wasting literature where similar mitochondrial features are seen was a study by Owen and colleagues in a mouse model of persistent skeletal muscle pathology following recovery from sepsis. Their study showed mitochondria with condensed cristae, mitochondria with displaced cristae, and mitochondria with a ruptured outer membrane (see Fig. 4 in that study (Owen *et al*., 2019)), similar to what we observed in response to Ca^2+^-induced mPT. Furthermore, a prior study by Fontes-Oliveira reported a wide variety of mitochondrial morphological alterations in cancer cachexia (Fontes-Oliveira *et al*., 2013) that also bear close resemblance to the impact of mPT seen in our experiments. On this basis, we suggest that in wasting conditions where these mitochondrial morphological alterations are present, mPT should be considered as a potential explanation.

### Inducing mPT in Skeletal Muscle Generates Phenotypes Common in Wasting Disorders

Skeletal muscle in wasting disorders often present with a common set of phenotypes including muscle atrophy (Friedrich *et al*., 2015; Martin *et al*., 2023), mitochondrial respiratory dysfunction (Protti *et al*., 2007; Neyroud *et al*., 2019; Kunz *et al*., 2022), and dismantling of the acetylcholine receptor (AChR) cluster at the neuromuscular junction (Liu *et al*., 2016; Sartori *et al*., 2021). Alterations in mitochondrial morphology (Fontes-Oliveira *et al*., 2013; Owen *et al*., 2019; Zhang *et al*., 2023) and reduced mitochondrial content (Brown *et al*., 2017; Neyroud *et al*., 2019) accompany these muscle changes, providing a compelling argument for considering the role of the mitochondrion in driving these pathological features. To test this, we performed a series of experiments to determine if inducing mPT could generate skeletal muscle phenotypes that are common in wasting disorders. To induce mPT we used a compound called Bz423, which binds to the oligomycin sensitivity conferring protein (OSCP) in the ATP synthase (Johnson *et al*., 2005; Starke *et al*., 2018) at the same proline residue as CypD to promote opening of the mPTP (Giorgio *et al*., 2013). To help validate the impact of mPT, in a subset of experiments we also employed use of a plant-derived Ca^2+^ ionophore called ferutinin to induce mPT by promoting mitochondrial Ca^2+^ overload (Ilyich *et al*., 2018; Denieva *et al*., 2026). Notably, our results show that Bz423 promotes mPT by reducing the Ca^2+^ threshold for mPT, and that inhibiting mPT with the triazole compound TR002 prevents this effect of Bz423 (Fig 2A). We found that Bz423-induced mPT was sufficient to induce signals (i.e., ROS, Casp3) that activate atrophy pathways and cause dismantling of the AChR cluster at the muscle endplate. We also observed that Bz423 was sufficient to induce: atrophy (Skinner *et al*., 2021), a complex I-specific deficit in mitochondrial respiration, increased colocalization of lysosomes with mitochondria, and fragmentation of the AChR cluster at the muscle endplate. The rescue of these Bz423-induced effects by mPT inhibitors is consistent with the actions of Bz423 being mediated through mPT. To further substantiate that the impact of Bz423 was due to its actions on mPT, in a subset of experiments we used the Ca^2+^ ionophore ferutinin to induce mPT. First, we showed that ferutinin reduced mitochondrial calcein and mitochondrial membrane potential. The principle of this assay is that upon the addition of cobalt the calcein signal is quenched in the cytoplasm, whereas mitochondrial calcein persists as long as the mPTP remains closed because cobalt cannot cross the inner mitochondrial membrane (Martel *et al*., 2012). However, upon opening of the mPTP cobalt enters the mitochondrial matrix to quench the calcein signal. Hence, a reduction in mitochondrial calcein indicates that mPT has occurred. Furthermore, when a reduced mitochondrial calcein is accompanied by reduced mitochondrial membrane potential, it is indicative of prolonged high conductance mPT rather than brief flickers (Petronilli *et al*., 2001). Thus, our results with ferutinin confirm its impact in inducing prolonged mPTP opening (Ilyich *et al*., 2018; Denieva *et al*., 2026). We then showed that similar to what was observed with Bz423, ferutinin induced a complex I-specific reduction in respiration. Finally, like Bz423 and a model of disuse in our previous study (Skinner *et al*., 2021), ferutinin also induced myotube atrophy. Thus, the similarity of impact between Bz423 and ferutinin is consistent with mPT being involved. The significance of our findings is that, in combination with the mitochondrial morphological features in wasting conditions (Kouidi *et al*., 1998; Fontes-Oliveira *et al*., 2013; Owen *et al*., 2019; Zhang *et al*., 2023; Pierre *et al*., 2025) that mirror the consequences of Ca^2+^-induced mPT in our experiments, they suggest that consideration of mPT as a likely contributor to muscle pathology in wasting conditions is warranted.

It is relevant to point out that the muscle fiber atrophy seen in response to Bz423 was prevented not only by targeting mPT, but also by quenching signals that are widely considered to drive muscle atrophy: mitochondrial ROS (Dodd *et al*., 2010; Powers *et al*., 2011; Ahn *et al*., 2022) and Casp3 activity (Du *et al*., 2004; Wang *et al*., 2010; Silva *et al*., 2015). We also previously showed in a multi-day single muscle fiber incubation that mPT was necessary for muscle fiber atrophy and acted upstream of mitochondrial ROS and Casp3 (Skinner *et al*., 2021). It is also relevant that Casp3 activation is implicated in the muscle wasting seen in cachexia (Belizario *et al*., 2001). Finally, a complex I-specific defect consequent to mPT is seen in liver cells (Briston *et al*., 2017), and Wrogeman and colleagues reported in 1973 that skeletal muscle mitochondria from dystrophic hamsters exhibit a complex I-specific respiratory defect that was associated with very high mitochondrial Ca^2+^ accumulation (the primary trigger for mPT) (Wrogemann *et al*., 1973). Thus, our observation that mPT induced by either Bz423 or ferutinin markedly reduced respiration with complex I substrates but yielded a normal increase in response to a complex II substrate and direct electron transfer to complex IV (assessed with Bz423 treatment only), is consistent with prior studies addressing the impact of mPT on mitochondrial respiratory function. Notably, a complex I-specific respiratory defect has also been reported in cancer cachexia (Neyroud *et al*., 2019) and in mice that survive sepsis (Pierre *et al*., 2025). It is important to acknowledge, however, that we do not know how long a complex I-specific deficit would persist after mPT is resolved nor whether acute changes in response to mPT (e.g., high ROS) may generate prolonged alterations (e.g., oxidative modifications to complex I) that would allow a complex I deficit to persist after mPT has resolved. Concerning the increase in lysosomal colocalization with mitochondria following induction of mPT, collapse of mitochondrial membrane potential is both a consequence of mPT (Petronilli *et al*., 2001; Zamzami & Kroemer, 2004) and a trigger for mitochondrial removal by mitophagy (Narendra *et al*., 2008; Narendra *et al*., 2010). Thus, our results are consistent with mPT causing depletion of mitochondria, a phenotype that is common in muscle wasting conditions. Finally, our results showing that inducing mPT in single mouse skeletal muscle fibers causes fragmentation of the AChR cluster in a manner that depends upon Casp3, are consistent with evidence that Casp3 causes AChR dismantling at the endplate (Wang *et al*., 2014) and places mPT as an upstream mechanism by which this occurs. Collectively, our results highlight pleiotropic consequences of mPT in generating pathological phenotypes in skeletal muscle. Our results in no way rule out the involvement of other mechanisms as contributors to muscle pathology but hint at a mechanistic hub that may be common to many wasting conditions.

### Transcriptional Reprogramming with mPT is Consistent with the Pathological Phenotypes Observed

To identify downstream effectors involved in the pathological phenotypes generated by mPT we induced mPT with Bz423 in C2C12 myotubes and isolated RNA for sequencing at 12, 24 and 48 h post-treatment. The number of DEGs peaked at 24 h, where the largest changes involved upregulation of pathways involved with fatty acid/lipid metabolism, consistent with extensive lipid membrane remodeling with mPT. We also observed upregulation of 38 FoxO signaling related genes, consistent with the myotube atrophy observed with induction of mPT. We saw reduction in amino acid metabolic pathways, and reductions in pathways such as cell cycle, mitochondrial translation elongation, and oxidative phosphorylation. Although small in magnitude, there were changes in pathways consistent with muscle atrophy (*Fbxo32* and the transcription factor *Foxo3)*, neuromuscular junction alterations (reductions in *Chrne*, the adult AChR subunit that is lost with denervation (Witzemann *et al*., 1991) and *Fgfbp1* which is involved in neuromuscular junction maintenance and repair (Taetzsch *et al*., 2017)), impaired mitochondrial respiratory capacity (reduced levels of several genes encoding subunits of mitochondrial complexes involved with oxidative phosphorylation), and mitochondrial depletion (reduced levels of genes encoding microtubule proteins involved in autophagy/mitophagy).

### Rationale for Therapeutic Targeting of mPT in Cancer Cachexia

To examine the potential promotion of mPT in skeletal muscle in wasting conditions, we tested whether factors present in TCM from cachexia-inducing pancreatic KPC tumor cells impacted mPT. The rationale for considering cancer cachexia as a likely condition where mPT is involved is based upon the well-established mitochondrial impact in cancer cachexia (van der Ende *et al*., 2018), manifesting as impaired mitochondrial respiratory function (Julienne *et al*., 2012; Kunz *et al*., 2022) (including a complex I-specific deficit reported previously in a colon C26 model (Neyroud *et al*., 2019)), reduced mitochondrial content (Fermoselle *et al*., 2013; Tzika *et al*., 2013) and mitochondrial cristae disruption (White *et al*., 2012; Fontes-Oliveira *et al*., 2013; Zhang *et al*., 2023), which mirror changes we see in response to inducing mPT *in vitro*. Furthermore, numerous tumor-host factors implicated in muscle impairment in cancer cachexia (Ji *et al*., 2011; Zhang *et al*., 2018; Chen *et al*., 2020; Talbert & Guttridge, 2022) activate mPT-induced cell death pathways in non-muscle cell types but have not been examined in skeletal muscle. Although mPT has not been directly considered, there is evidence for Ca^2+^ dyshomeostasis (Dasgupta *et al*., 2025; Gand *et al*., 2025) and sarcoplasmic reticulum disruption (Fontes-Oliveira *et al*., 2013) in cancer cachexia (Ca^2+^ is the primary trigger for mPT), and a recent study reported upregulation of transcriptional pathways related to apoptosis, mitochondrial swelling, and mitochondrial permeabilization in cachectic muscle (Gicquel *et al*., 2024). To directly test if mPT occurs in response to cachectic tumor-derived factors, we exposed mouse C2C12 myoblasts to KPC TCM in a Calcein-Cobalt assay. KPC TCM reduced mitochondrial Calcein localization and reduced mitochondrial membrane potential (reduced TMRM fluorescence) (Fig 8A & B), showing that KPC TCM induces long-lasting mPT in muscle cells.

The Ca^2+^ threshold for mPT is regulated by the peptidyl-prolyl *cis-trans* isomerase CypD, and this threshold is reduced in pathological conditions linked to mPT such as ischemia-reperfusion injury (Murphy, 2022). To reveal mechanisms that might promote mPT in skeletal muscle in cancer we tested whether exposure to TCM from pancreatic KPC cells would reduce the Ca^2+^ threshold for mPT in mouse skeletal muscle and determined its dependence on CypD. Our data show that KPC TCM markedly reduced the Ca^2+^ threshold for mPT but that this impact was ablated by CypD-dependent mPT inhibition using Alisporivir. Furthermore, skeletal muscle CypD knockout prevented 75% of the reduction in Ca^2+^ threshold induced by tumor-conditioned media. That this occurred with only a 60 min incubation in KPC TCM and was significantly attenuated by CypD knockout is consistent with modulation of mPT occurring secondary to post-translational modifications to CypD (e.g., S191 phosphorylation of CypD reduces the Ca^2+^ threshold for mPT (Hurst *et al*., 2020)), since post-translational modification events like phosphorylation can occur in response to a stimulus on the order of seconds (Gelens & Saurin, 2018). Our results do not rule out additional mechanisms contributing to an increased burden of mPT in skeletal muscle with cancer cachexia, such as impaired Ca^2+^ handling (Gand *et al*., 2025), sarcoplasmic reticulum dysfunction (Fontes-Oliveira *et al*., 2013) and histamine-dependent Ca^2+^ release (Dasgupta *et al*., 2025) that increase mitochondrial Ca^2+^ exposure in cancer cachexia. Regardless of these possibilities, our results show that factors released from cachexia-inducing tumor cells promote mPT in skeletal muscle cells rapidly, in part by reducing the Ca^2+^ threshold for mPT in a manner that was significantly attenuated by CypD knockout.

To consider the potential burden of mPT *in vivo* with cancer cachexia we examined the degree of overlap in DEGs between mPT in C2C12 myotubes and publicly available time-course transcriptomic data of diaphragm muscle from mice with pancreatic cancer (Neyroud *et al*., 2025). Consistent with mPT occurring in cancer cachexia, there was a progressive increase in DEGs common to mPT and cancer cachexia that was amplified during the period when muscle wasting occurred. Examples of overlapping DEGs with mPT and cachexia included a common downregulation of pathways involved with mitochondria (tricarboxylic acid cycle, fatty acid metabolism) during the cachectic period. Furthermore, key commonly upregulated pathways included cell death, denervation/neuromuscular junction genes and cholesterol biosynthesis. Our analysis of common DEGs revealed substantial overlap in these pathways not only in the same direction but also in the specific genes involved. Although *a priori* we expected some overlap, we were surprised at the high magnitude of concordance, particularly during the cachectic phase, given that the mPT transcriptome was generated using *in vitro* cultured myotubes whereas the cancer cachexia transcriptome data came from diaphragm muscle of an *in vivo* mouse model of orthotopic pancreatic cancer cachexia. On the other hand, there were notable differences as well in that the *in vivo* cachexia model exhibited upregulation of pathways involved in muscle fibrosis that did not overlap with DEGs seen with mPT. This likely reflects the contribution of fibroadipogenic progenitor cells and infiltrating inflammatory cells in the *in vivo* model that would not be seen in our C2C12 cell culture conditions. On the basis of this comparison, it appears highly likely that mPT is occurring in cancer cachexia during the period where muscle pathology occurs and contributes to muscle pathology. The strong inverse relationship between diaphragm diameter and the number of mPT DEGs in the diaphragm transcriptome as cachexia progressed supports this idea. To more directly test the impact of tumor-derived factors in driving atrophy in a manner that depends upon mPT, we determined if mPT inhibition could reduce the atrophy induced by KPC TCM in both C2C12 and human primary myotubes. Consistent with an important role of mPT in driving atrophy by tumor-derived factors, mPT inhibition attenuated (C2C12 myotubes) or completely prevented (human primary myotubes) atrophy induced by KPC TCM.

## Conclusions

We conclude that mPT is sufficient to recapitulate multiple features of mitochondrial morphological disruption and muscle pathology observed in wasting conditions and appears to operate upstream of established pathways involved in atrophy, neuromuscular junction maintenance, mitochondrial oxidative phosphorylation and mitophagy. Furthermore, tumor-derived factors increase mPT in muscle cells, due in part to a reduced Ca^2+^ threshold for mPT that was significantly attenuated by knockout of the mPT-modulating mitochondrial protein, CypD. We found significant overlap in DEGs induced by mPT and pancreatic cancer cachexia that was particularly prevalent during the period of muscle pathology and closely tracked the progression of muscle atrophy. Finally, we found that mPT inhibition attenuated atrophy induced by KPC TCM in both C2C12 and human primary myotubes, demonstrating the importance of mPT in driving atrophy by tumor-derived factors. Our findings support the idea that mPT may be a core mechanism of muscle pathology in wasting conditions, and that therapeutic targeting of mPT is worthy of study as a strategy to preserve skeletal muscle mass and function in wasting conditions.

## Additional Information Section

### Data Availability

Complete pathway analysis for the mPT transcriptome can be found in Supplemental Dataset 2. Raw sequencing data can be found at the GEO accession, Series GSE317195. Raw data for other analyses is available on request.

### Author Contributions

MS, CL, SS, MRV, SGN, MP, and AGM were involved with data acquisition, analysis and interpretation. RTH, TER, MSC, and DWW were involved with the conception and design of the work. All authors contributed to drafting the work and revising it for intellectual content. All authors have approved the final version, agree to be accountable for all aspects of the work, all persons designated as authors qualify as authors, and all qualified for authorship are listed.

Artificial Intelligence Generated Content: No generative AI tools were used for any aspect of this manuscript.

This work was supported by NIAMS R21AR084591, NIA R56AG066758, and a University of Florida PHHP Research Innovation Fund award. M. Semel was supported by a predoctoral fellowship from the BREATHE T32 (T32HL134621), and C. Lukasiewicz was supported by a University of Florida Graduate Student Opportunity Award.

## Supporting Information

We include 6 figures in the appendix containing data that includes validation of the mkoPPIF mouse model, validation of that ferutinin induces long-lasting mPT events, shows the impact of a higher dose of ferutinin on mitochondrial respiration, shows the impact of mPT on state II respiration when using Bz423 or ferutinin, a second validation experiment demonstrating the impact of Bz423-induced mPT on co-localization of lysosomes with mitochondria in C2C12 myotubes, and heatmaps detailing specific DEGs induced by mPT that are relevant to the muscle pathological phenotypes of atrophy, integrity of AChR cluster, mitochondrial oxphos, and autophagy/mitophagy.

## Appendix

**Fig A1.**
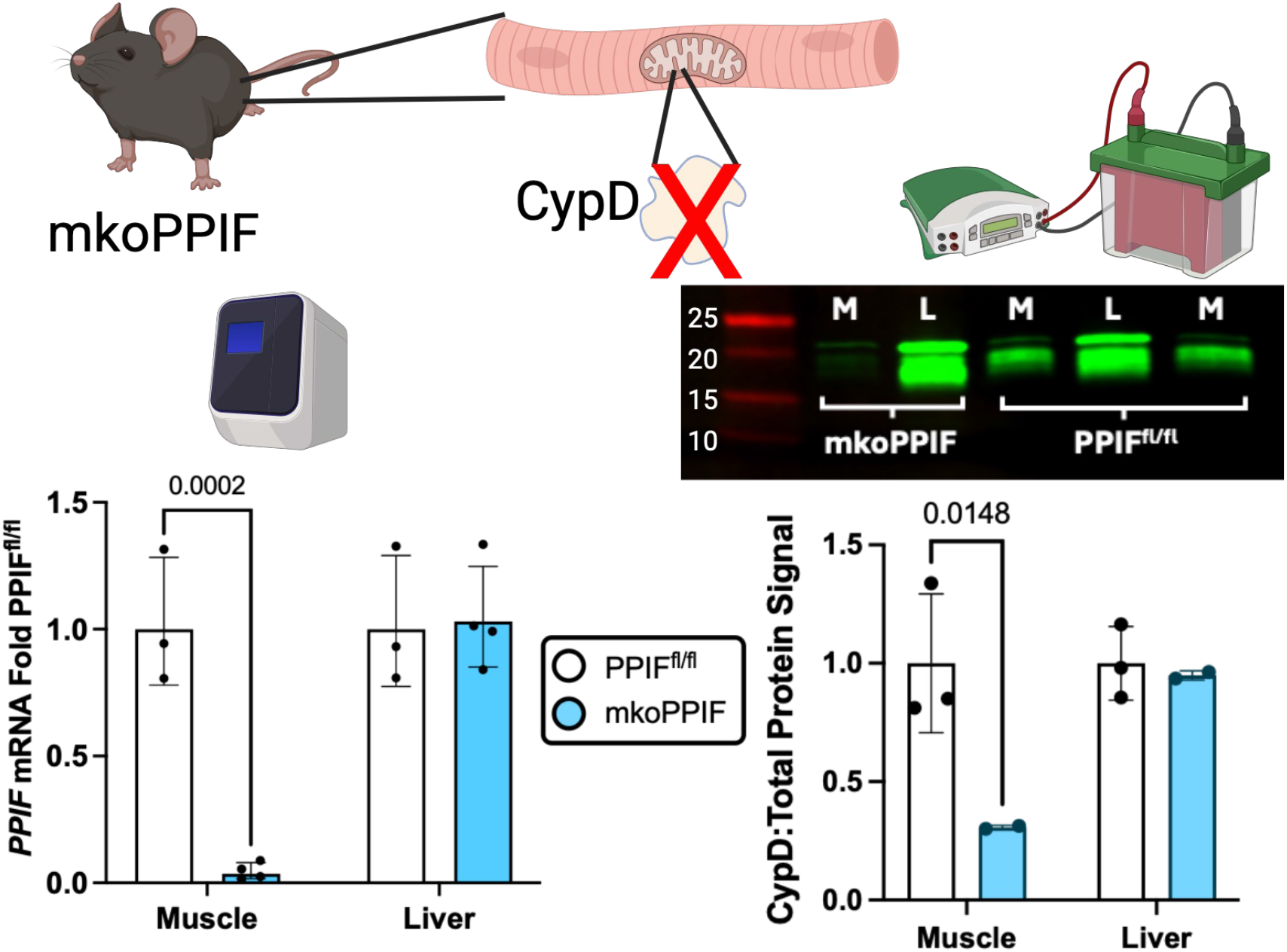
Validation of muscle-specific CypD knockout (mkoPPIF) mouse. One-way ANOVA with Sidak post-hoc test.

**Fig A2.**
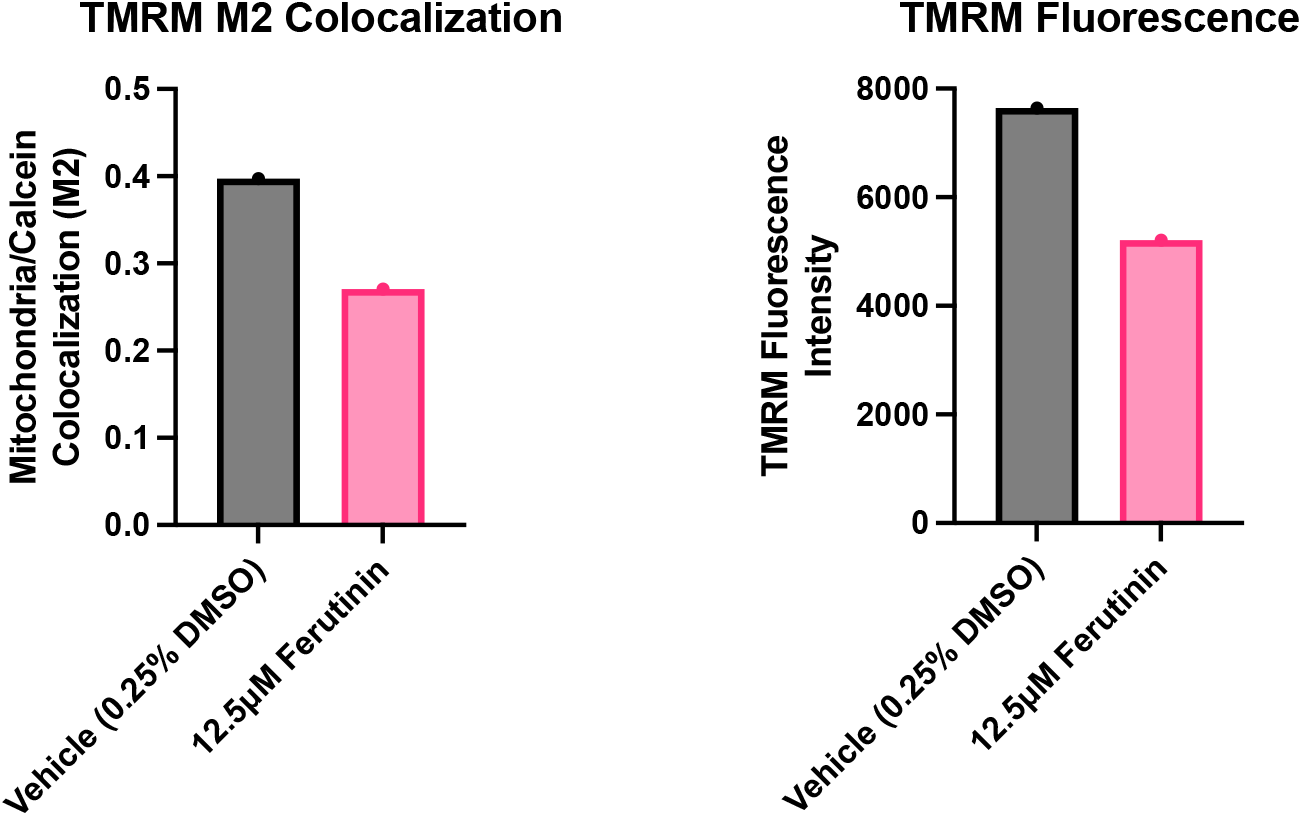
C2C12 myoblasts were incubated with 12.5 μM ferutinin for 30 min before undergoing a Calcein-Cobalt assay. Ferutinin induced a reduction in mitochondrial Calcein and mitochondrial membrane potential (TMRM), showing that ferutinin generates long-lasting mPT events.

**Fig A3.**
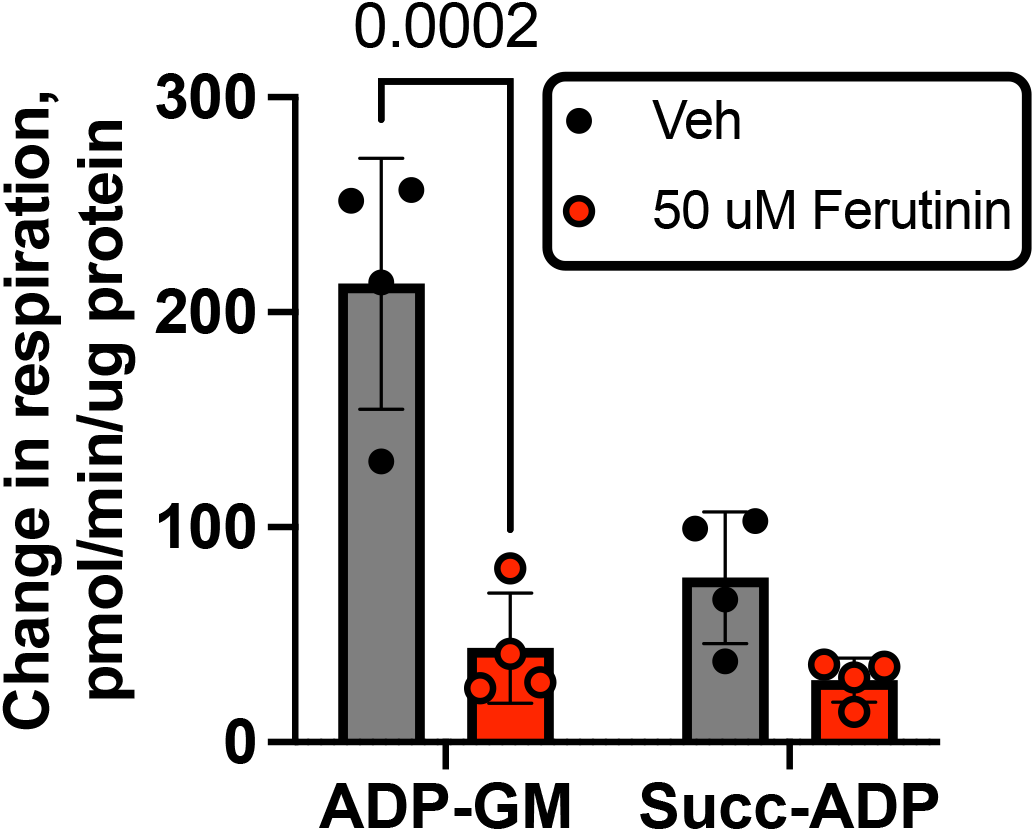
Treating isolated skeletal muscle mitochondria with 50 μM ferutinin (Ca^2+^ ionophore) causes a complex I-dominant decline in maximal respiratory capacity. One-way ANOVA with Sidak post-hoc test.

**Fig A4.**
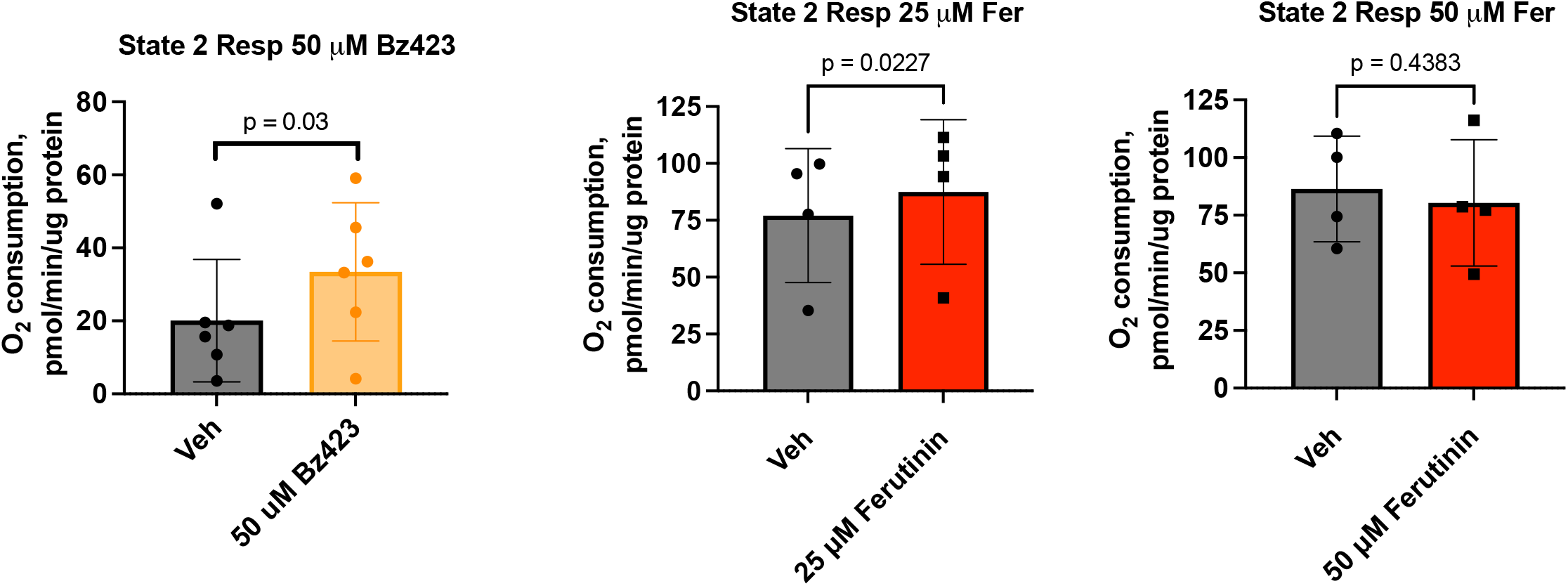
Inducing mPT with either 50 μM Bz423 (P=0.03) or 25 μM ferutinin (P=0.023) increases state II respiration in isolated skeletal muscle mitochondria. This was not seen with 50 μM ferutinin. T-test.

**Fig A5.**
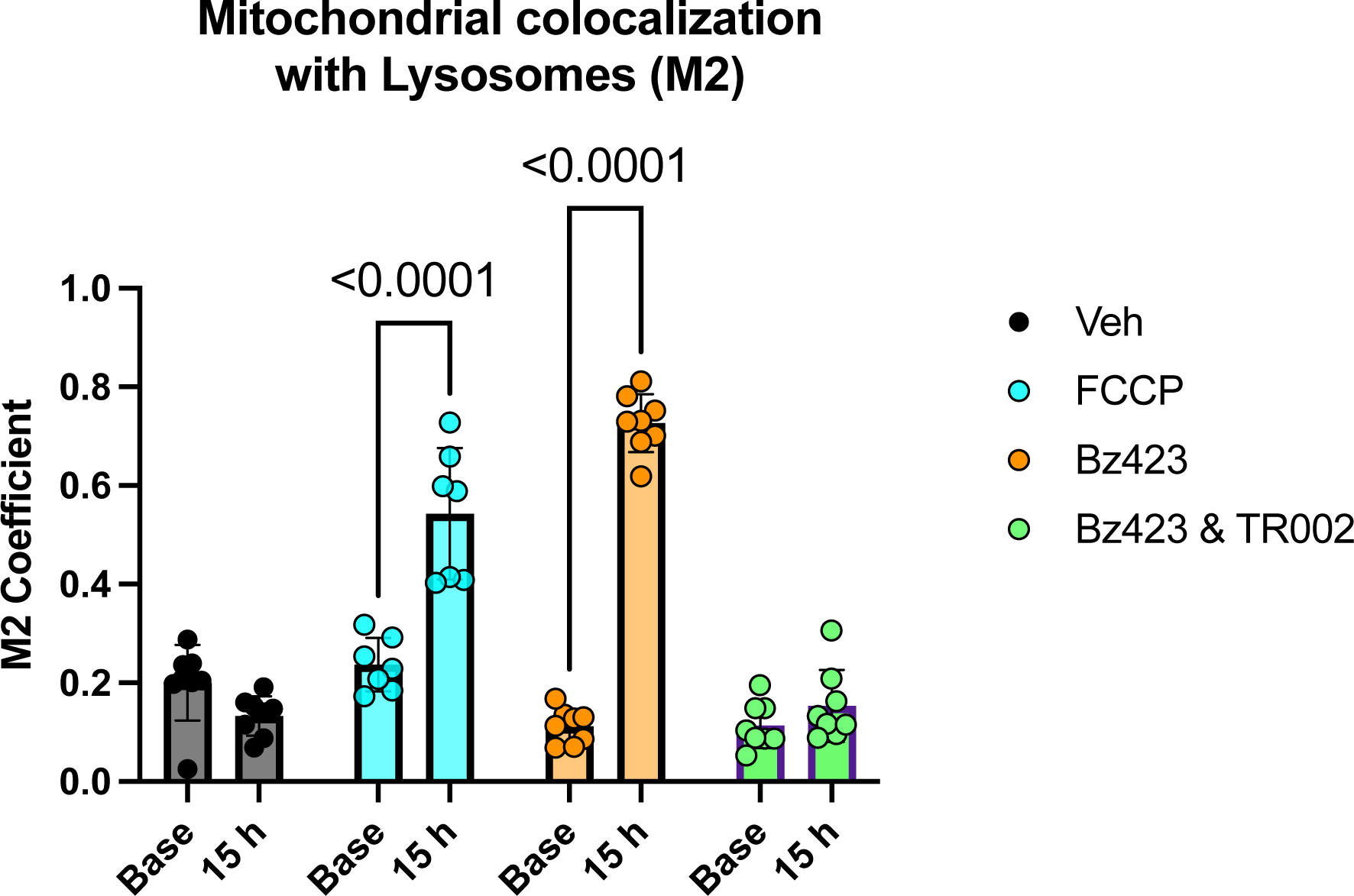
Repeat technical replicate of experiment showing that Bz423-induced mPT increases lysosomal colocalization with mitochondria. One-way ANOVA with Sidak post-hoc test.

**Fig A6.**
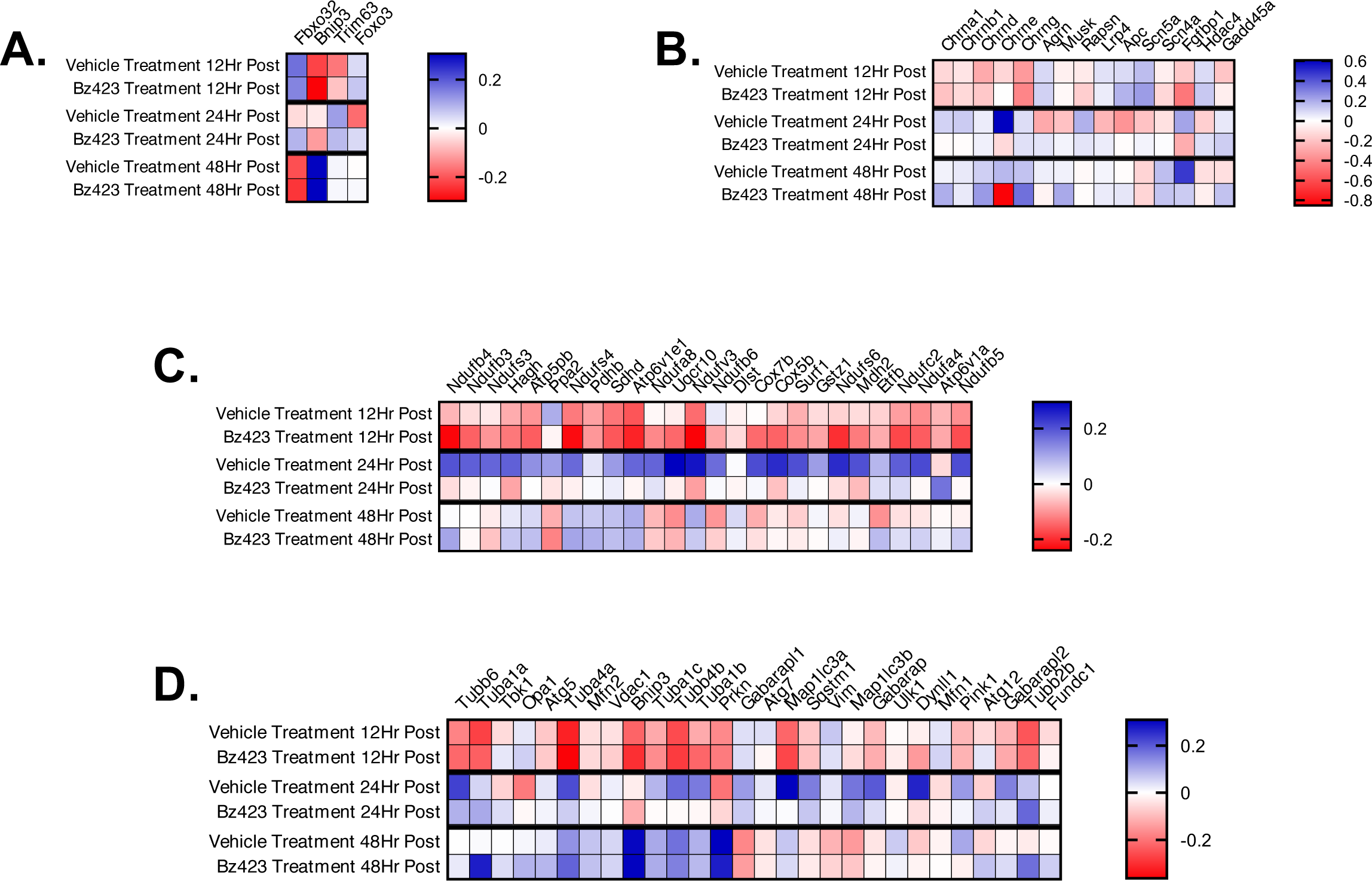
Inducing mPT in C2C12 myotubes alters pathways relevant to diverse muscle pathological phenotypes. Heatmaps showing mPT-induced pathway-specific changes for transcripts related to A: atrophy, B: integrity of AChR cluster, C: mitochondrial oxphos, and D: autophagy and mitophagy.

